# Prediction of symptomatic and asymptomatic bacteriuria in spinal cord injury patients using machine learning

**DOI:** 10.1101/2024.08.09.607254

**Authors:** M. Mozammel Hoque, Parisa Noorian, Gustavo Espinoza-Vergara, Joyce To, Dominic Leo, Priyadarshini Chari, Gerard Weber, Julie Pryor, Iain G. Duggin, Bonsan B. Lee, Scott A. Rice, Diane McDougald

**Affiliations:** Australian Institute for Microbiology & Infection, University of Technology Sydney, Sydney, NSW, Australia; Spinal cord injury unit, Royal North Shore Hospital, NSW, Australia; Royal Rehab Group, Sydney, NSW, Australia; Susan Wakil School of Nursing and Midwifery, University of Sydney, Sydney, Australia; Department of Spinal and Rehabilitation Medicine, Prince of Wales Hospital, Sydney, NSW, Australia; Neuroscience Research Australia (NEURA), Sydney, NSW, Australia; CSIRO, Microbiomes for One Systems Health, Agriculture & Food, Westmead NSW

**Keywords:** Spinal cord injury, Bacteriuria, Urinary tract infections, 16S rRNA, Catheter, Urine, Machine learning, Prediction

## Abstract

**Background:** Individuals with spinal cord injuries (SCI) frequently rely on urinary catheters to drain urine from the bladder, making them susceptible to asymptomatic and symptomatic catheter-associated bacteriuria and urinary tract infections (UTI). Proper identification of these conditions lacks precision, leading to inappropriate antibiotic use which promotes selection for drug-resistant bacteria. Since infection often leads to dysbiosis in the microbiome and correlates with health status, this study aimed to develop a machine learning-based diagnostic framework to predict potential UTI by monitoring urine and/or catheter microbiome data, thereby minimising unnecessary antibiotic use and improving patient health.

**Results:** Microbial communities in 609 samples (309 catheter and 300 urine) with asymptomatic and symptomatic bacteriuria status were analysed using 16S rRNA gene sequencing from 27 participants over 18 months. Microbial community compositions were significantly different between asymptomatic and symptomatic bacteriuria, suggesting microbial community signatures have potential application as a diagnostic tool. A significant decrease in local (alpha) diversity was noted in symptomatic bacteriuria compared to the asymptomatic bacteriuria (*P* < 0.01). Beta diversity measured in weighted unifrac also showed a significant difference (*P* < 0.05) between groups. Supervised machine learning models trained on amplicon sequence variant (ASVs) counts and bacterial taxonomic abundances (Taxa) to classify symptomatic and asymptomatic bacteriuria with a 10-fold cross-validation approach. Combining urine and catheter microbiome data improved the model performance during cross-validation, yielding a mean area under the receiver operating characteristic curve (AUROC) of 0.91-0.98 (Interquartile range, IQR 0.93-0.96) and 0.78-0.91 (IQR 0.86-0.88) for ASVs and taxonomic features, respectively. ASVs and taxa features achieve a mean AUROC of 0.85-1 (IQR 0.93-0.98) and 0.69-0.99 (IQR 0.78-0.88) in the independent held-out test set, respectively, signifying their potential in differentiating symptomatic and asymptomatic bacteriuria states.

**Conclusions:** Our findings demonstrate that signatures within catheter and urine microbiota could serve as tools to monitor the health status of SCI patients. Establishing an early warning system based on these microbial signatures could equip physicians with alternative management strategies, potentially reducing UTI episodes and associated hospital costs, thus significantly improving patient quality of life while mitigating the impact of drug-resistant UTI.

## Background

Spinal cord injury patients are at high risk of catheter-associated urinary tract infection (CAUTI) [1]. CAUTI typically manifests as a bacteriuria which is generally defined as a urine culture with at least 10^8 colony forming units (CFU)/L, of an identified microorganism(s) [2]. Bacteriuria can be further classified as asymptomatic bacteriuria (AB) or symptomatic bacteriuria (SB) [3]. Patients with AB generally lack signs and symptoms of UTI and do not require treatment [4, 5]. Conversely, SB is associated with symptoms such as fever, urethral and bladder inflammation, and potential renal scarring among other symptoms [6]. While bacteriuria is a significant risk factor for CAUTI in SCI patients, differentiating AB from SB can be challenging due to limitations in diagnosis and the influence of various etiologic factors [7, 8]. Accurate identification of these conditions is crucial because CAUT often necessitates extensive antibiotic use. Unfortunately, this therapy is becoming less effective due to the emergence of multidrug-resistant (MDR) bacteria. This poses life-threatening risks and creates a significant economic burden on public health systems. Healthcare costs associated with CAUTI are rising, with estimates suggesting millions of dollars are spent annually on treating hospitalisations caused by CAUTI, resulting in over 19,000 deaths in the United States [9–11].

The main cause for the development of CAUTI has been associated with the presence of pathogenic bacteria in microbial biofilms formed on the surface of urinary catheters. Reports show that CAUTI are predominantly caused by *Escherichia coli*, *Proteus mirabilis*, *Pseudomonas aeruginosa, Klebsiella pneumoniae* and *Enterococcus faecalis*, bacteria that have been associated with biofilm formation in urinary catheters [3, 12, 13]. Routinely, the diagnosis of CAUTI involves pathogen identification by urine screening followed by antibiotic treatment based on antibiogram reports produced by pathology laboratories. Despite the availability of screening and diagnostic tools for CAUTI, there is a lack of strategies that can successfully predict CAUTI. However, advances in the study of catheter-associated biofilm communities by next generation sequencing technologies can generate valuable information to build predictive platforms for CAUTI. For example, reports based on sequence analysis of urinary catheter associated-biofilms have shown that microbial communities associated with urinary catheters from long-term catheterised patients are diverse and show variability after UTI events or antibiotic treatments [14–16]. This fact leads to the idea that critical changes in the microbial community composition of urinary catheters can reveal the early steps of a UTI event. Thus, detection of early shifts in biofilm communities can be explored to establish ‘community thresholds’ which might serve as an early warning approach before a UTI event.

Machine learning approaches have been implemented for the prediction of various disease states [17]. Machine learning’s ability to capture subtle differences in feature abundances using 16S rRNA data allows for accurate prediction and classification of diseases [18–21]. Previous studies have explored machine learning models to predict UTI based on patient demographic information, biochemical and immunological markers [22–25]. However, demographic data alone could not explain the underlying causes of the infection and none of the studies underwent precise characterisation. Using a machine learning-based diagnostic framework, this work aimed to investigate a predictive platform for CAUTI based on microbial communities in the urine and catheters of patients.

Here, twenty-seven SCI participants were monitored longitudinally for changes in their urinary and catheter microbiome. We hypothesised that monitoring changes in the bacterial communities colonising patients’ catheters and urine can serve as an early warning system for impending CAUTI. Our data suggests that the composition of these microbial communities is dynamic, shifting in response to factors such as antibiotic use or pathogen colonisation. The results reveal that microbial signatures within urine and catheter could be used as a potential predictor of asymptomatic and symptomatic bacteriuria with high accuracy using a supervised machine learning model. This early diagnostic tool could empower clinicians to intervene before a full-blown infection develops. This approach has the potential to improve patient outcomes, reduce healthcare costs, and mitigate the spread of MDR pathogens.

## Methods and materials

### Study cohort and baseline characteristics

Participants were recruited from four specialist SCI units (Prince of Wales Hospital, Royal North Shore Hospital, Royal Rehab Ryde and Fairfield West Medical Centre) in New South Wales, Australia. They were all adults, aged 18 years and older with inclusion criteria of stable SCI, stable neurogenic bladder management for at least 4 weeks and agreement to fortnightly telephone consultations over 18 months. They also agreed to have their urine and catheter specimens and extracted DNA from these specimens stored for future studies. Exclusion criteria included long-term antibiotic therapy, immunosuppressant use, invasive mechanical ventilation, chronic infections, surgical bladder interventions, severe renal/hepatic failure, and concurrent enrolment in any intervention studies.

The samples were collected during regular catheter changes by their care team. Only samples that would be otherwise discarded were collected, including 5 cm of the bladder end of the used catheter and urine samples from the new catheter. Participants reporting potential UTI symptoms were instructed to contact their medical practitioner, providing additional urine samples and 5 cm of the old catheter to the research team if changed. Baseline and first UTI event samples underwent routine pathology tests. Diagnosis of symptomatic bacteriuria/ UTI relied on subjective complaints and lab findings, following diagnostic criteria first published in the SINBA randomised controlled trial [26–28]. It was crucial to distinguish new or increased symptoms from chronic issues, as many symptoms alone did not justify treatment. UTI symptoms were defined by new onset symptoms and laboratory evidence of UTI.

Between November 2021 and March 2023, 39 potential participants expressing interest were screened, with 27 enrolled. Participants were predominately male (66.7%), female (33.3%) with a mean age of 55 years (Supplementary Table S1). Most used suprapubic catheters, with only one participant using an indwelling urinary catheter. The median time since SCI was 18 years (range: 98 days to 56 years). The research team received subjective complaints of CAUTI symptoms, designating 67 UTI/ symptomatic bacteriuria events based on symptoms, pathology analysis and self-reported data

### Sample processing

Samples were processed according to previously described methods with some modifications [14]. A 5 cm section of the bladder end of the used catheter was collected in a sterile container with 5 mL of sterile saline (0.9% NaCl). A fresh catch urine specimen was also collected from the newly installed catheter in a separate sterile container. Both specimens were transported to the laboratory by courier or overnight express post and processed immediately on the day of receiving samples. The catheter was cut into half lengthwise along the inflation line using a sterile scalpel blade. The content inside was soaked with 1 mL of saline. A 1 mL syringe plunger was used to dislodge the soaked content by running the plunger back and forth on the two halves of the catheter pieces. The catheter pieces and plunger were transferred to the original container with saline, vortexed and sonicated in an ultrasonic water bath (Powersonic 420, Thermoline Scientific) at medium power for 1 min, after which time 1 mL of catheter cell suspension as well as 12 mL of urine samples were centrifuged at 5000 g for 5 min. The supernatants discarded from both tubes and the cell pellets were stored at −20°C for DNA extraction. The remaining catheter cell suspensions were centrifuged at 5000 g for 5 min and cell pellets were frozen at −80°C in glycerol for future analysis.

### DNA extraction

Total DNA was extracted from catheter and urine pellets obtained in the previous step using DNeasy PowerSoil Pro Kit (Qiagen, cat. no. 47016) according to the manufacturer’s instructions except that the final elution was in 50 µL of water. The quality and quantity of the isolated DNA were determined using a NanoDrop spectrophotometer (Thermo Fisher Scientific, USA). The DNA samples were stored at −20°C before further analysis.

### Library preparation and amplicon sequencing

The V4 region of the bacterial 16S rRNA gene was amplified for sequencing by a two-stage PCR process. The first PCR was carried out using 10 ng of genomic DNA using 515F (5′-GTGYCAGCMGCCGCGGTAA 3′) and 806R (5′-GGACTACNVGGGTWTCTAAT 3′) primers including the Illumina adapters and KAPA HiFi HotStart ReadyMix (Cat No.KR0370-v14.22; Roche, Switzerland) under the PCR cycle: initial denaturation at 95°C for 2 min; followed by 20 cycles of: 95°C for 15 s, 60°C for 15 s, 72°C for 30 s; and a final extension step at 72°C for 1 min and hold at 4°C. The PCR products from the first PCR were diluted in water (1:40) and 3 µL of the diluted products were used as templates for the second PCR to add unique barcodes to each sample. PCR conditions were the same as the first PCR except that only 10 cycles were used. Two µL of each final product was pooled into one tube and solid-phase reversible immobilisation (SPRI) beads (Beckman Coulter, USA) were used to remove excess primers. The cleaned libraries were sequenced on an Illumina MiSeq v2 Nano 2 x 150 bp to assess read counts. The final normalised libraries were sequenced on an Illumina MiSeq v3 2 x 300 bp.

### Sequence processing and alignment

Sequencing data were analysed using the Quantitative Insights Into Microbial Ecology 2 program (QIIME2) version 2022.8.0 [29]. Raw fastq sequencing reads were imported to QIIME2 in iHPC environment using qiime ‘tools import’ plugin. The reads were filtered to remove sequencing primers using cutadapt [30]. The primer-trimmed sequences were denoised and clustered to amplicon sequence variants (ASVs) using DADA2 plugins. At the denoising step, the forward and reverse reads were truncated at position 220 and 160 bp, respectively, to retain only high-quality sequences [31]. The orientation of the denoised sequences was corrected by aligning to the reference sequences using ‘rescript orient-seq’ plugins [32]. For taxonomic assignments, the oriented sequences were aligned to the greengenes2 16S rRNA reference database (V4 region) using ‘feature-classifier’ plugin [33]. The Naive Bayes classifier pre-trained on the V4 region was obtained from greengenes2 data repository. Greengenes2, released in 2023, is the latest 16S rRNA reference database and contains high-quality full length 16S sequences from the Living Tree Project with updated taxonomic information. The feature table was filtered to remove unassigned features based on the taxa table obtained during the alignment step. The feature table was further subjected to taxonomic filtering to remove low abundant phyla (if feature frequency less than 5). The filtered sequences were aligned using ‘mafft’ and ‘fasttree’ plugins to generate rooted phylogenetic trees [34, 35].

### Microbiome diversity and taxonomic analysis

The feature table, taxonomic table and rooted tree obtained from the previous section were imported to build phyloseq object in R program using phyloseq package [36]. All samples were rarefied at 11,000 reads with the phyloseq function ‘rarefy_even_depth’ to normalise the variance [37]. Rarefaction retained 591 samples (307 catheter and 284 urine) and 18 of the samples were excluded from the analysis due to insufficient reads. All the downstream alpha diversity, beta diversity, taxonomic and machine learning analyses were performed with the rarefied dataset. The alpha diversity ‘Shannon index’ was computed using the R package ‘Microbiome’ [38]. Principal coordinates analysis (PCoA) was carried out on the beta diversity (weighted and unweighted unifrac) distance metrics using ‘microeco’ R package [39]. Taxonomic abundances were calculated using ‘microeco’ package at different taxonomic level. Taxonomic abundances data for machine learning were prepared using the ‘trans_classifier’ function of the microeco package in R. Shared and unique taxa analyses were also conducted using the ‘microeco’ package. Data were visualised using the ‘ggplot2’ package in R version 4.2 [40].

### Supervised machine learning

The raw ASVs counts and taxonomic abundance data were pre-processed in three steps using the R package mikropml [41]. First, the raw ASVs counts and taxonomic abundance data were pre-processed using the default method. Briefly, the default method normalised the data by centering and scaling and removed variables with near-zero variance. Second, the unique ASVs or Taxa belonging to AB and SB were also subjected to mikropml pre-processing to remove zero variance features only. In the third and final step, the feature lists from the first and second steps were combined and again subjected to pre-processing using mikropml to remove zero variance features. The pre-processed data were utilised to initiate the supervised machine learning pipeline using the PyCaret package (version 3.2) in python with default parameters unless otherwise stated [42]. For 20 random seeds, transformed data were subjected to stratified (proportional class distribution) split to obtain 80% training and 20% held-out sets. We used 10 iterations of stratified 10-fold cross-validations to ensure the robustness of our approach and to precisely evaluate the prediction power of the models. We applied SMOTE (Synthetic Minority Over-sampling Technique) to fix imbalances in the distribution of the target class in the training set during the PyCaret setup function. We also removed outliers using sklearn’s “IsolationForest” method with default threshold (0.05) during the setup function.

A second round of feature selection was applied to remove additional features based on the classic feature section method within PyCaret setup with the ‘n_features_to_select’ parameter was set at 0.9. During model optimisation, a total of 16 machine learning algorithms from the scikit-learn library were used to construct initial models (Supplementary Table S2) [43]. The top three models, based on balanced accuracy, were blended and tuned. The blended and tuned model performance was evaluated on both cross-validation and held-out sets. We evaluated model performance based on several metrics including AUROC (summarises trade-off between sensitivity and specificity across all possible thresholds) and AUPRC (focuses on the trade-off between precision and recall). In addition to these two metrics, we also provided accuracy (correct prediction / all prediction), precision (true positives divided by the total number of positive predictions), recall (weighted average of sensitivity and specificity), balanced accuracy (arithmetic average of sensitivity and specificity) and F1 scores (harmonic mean of the precision and recall) (Supplementary Tables S3 and S4). All hyperparameters were automatically tuned and optimised by the PyCaret. Finally, the most important ASVs and Taxa contributing to model performance were determined by the feature importance score extracted from the top performing model.

### Statistical analysis

Statistical significance for the alpha diversity (Shannon index) metric was calculated with non-parametric Wilcoxon test in R. Statistical significance for beta diversity (weighted and unweighted unifrac distance) metrics were determined by Permutational Multivariate Analysis of Variance (PERMANOVA) with a number of 999 permutations using QIIME2 ‘diversity beta group significance’ plugin [44]. Differences in AUROC scores between cross-validation and held-out set were determined by no-parametric Wilcoxon test in R. Differences in ASVs and taxa abundances were also determined by no-parametric Wilcoxon test in R.

## Results

### Evaluation of microbial community composition in asymptomatic and symptomatic bacteriuria

To investigate the microbial community composition in asymptomatic (AB) and symptomatic (SB) bacteriuria, we performed 16S rRNA sequencing analysis on a total of 300 urine and 309 catheter samples collected from 27 participants (Fig. 1a). A total of 35,101,926 high quality sequences were produced, with a median of 61,613 sequences per sample. After quality filtering, sequencing reads were clustered into 1246 amplicon sequence variants (ASVs) of which 1128 ASVs remained after rarefaction. The AB group harboured more distinct ASVs compared to the SB group (Fig. 1b). Out of 1128 ASVs, 874 (77%) and 79 (7%) were unique to AB and SB, respectively, with 175 (16%) shared by both groups. The identified ASVs belonged to diverse phylogenetic lineages, spanning 8 phyla, 11 classes, 43 orders, 68 families, 138 genera and 144 species (Fig. 1c). Nearly all (mean ∼100%) ASVs were classified to the family level, while a mean of 76.1% and 34.1% were assigned to genus and species levels, respectively. The majority of the ASVs were Proteobacteria (n=433, 38.4%), followed by Firmicutes_D (n=183, 16.2%), Actinobacteriota (165, n=14.6%), Firmicutes_A (n=112, 9.9%), Firmicutes_C (n=98, 8.7%), Bacteriodota (n=85, 7.5%), Fusobacteriota (n=35, 3.1%) and Campylobacterota (n=17, 1.5%). Enterobacteriaceae (n=282, 25%) and Pseudomonadaceae (n=70, 6%) were the largest contributors to the Proteobacteria phyla (Supplementary Fig. S1).

**Fig. 1.**
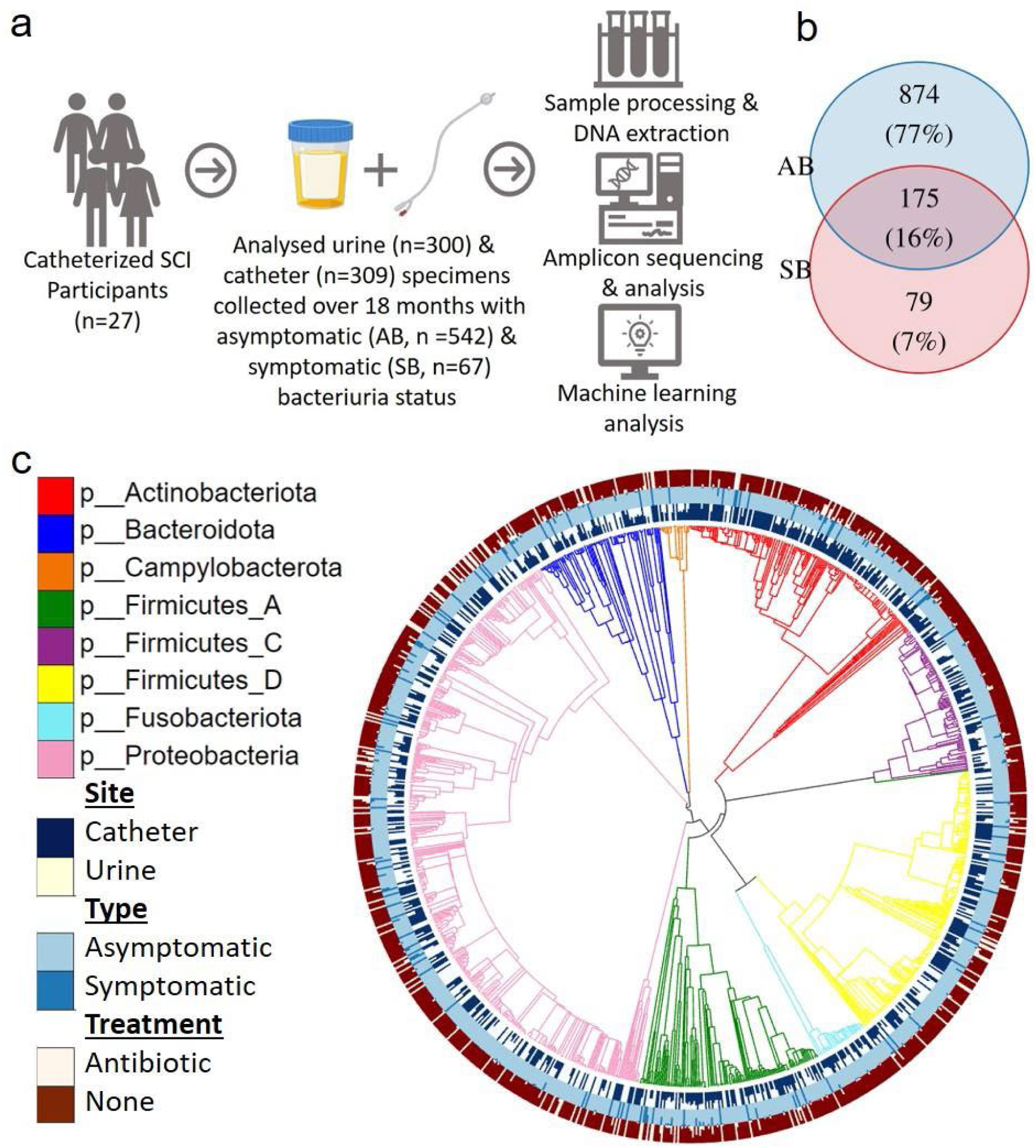
Study design and distribution of 16S rRNA amplicon sequence variants. (a) The schematic shows the overview of the study design and workflow. (b) Venn diagram showing the unique and shared ASVs between asymptomatic (AB) and symptomatic (SB) bacteriuria groups. (c) Phylogenetic tree for 1128 microbiome members constructed based on 16S rRNA amplicon sequence variants. Tree branches are coloured based on their respective phylum. The inner, middle and outer bar plot rings indicate the proportion of counts split by site (catheter vs urine), type (AB vs SB) and antibiotic use (with and without), respectively, as indicated in the legend.

We assessed and compared the microbial community composition between AB and SB groups for urine and catheter samples separately as well as when combined. Alpha diversity, measured by Shannon index (accounting for both species abundance and evenness), was significantly lower in SB compared to AB, both in the combined and individual datasets (Fig. 2a-c). The mean Shannon index was 1.3 for AB (IQR 0.8-1.8) compared to 1.0 for SB (IQR 0.4-1.4) in the combined dataset. Beta diversity analysis using weighted unifrac distances revealed significant differences in community composition between AB and SB groups on the combined dataset (PERMANOVA, Pseudo-F = 2.5, *P* = 0.02) (Fig. 2d). While these differences were not statistically significant when analysed separately for urine or catheter samples, a moderate difference was observed for urine (*P* = 0.1) when compared to catheter (*P* = 0.2) (Fig. 2e-f). These findings suggest an altered community composition in SCI patients with symptomatic bacteriuria. Participant-specific analysis showed distinct clustering of microbial communities between AB and SB groups in participants one, six, eight, nine, eleven, twenty-five and twenty-six (Supplementary Fig. S2 and S3).

**Fig. 2.**
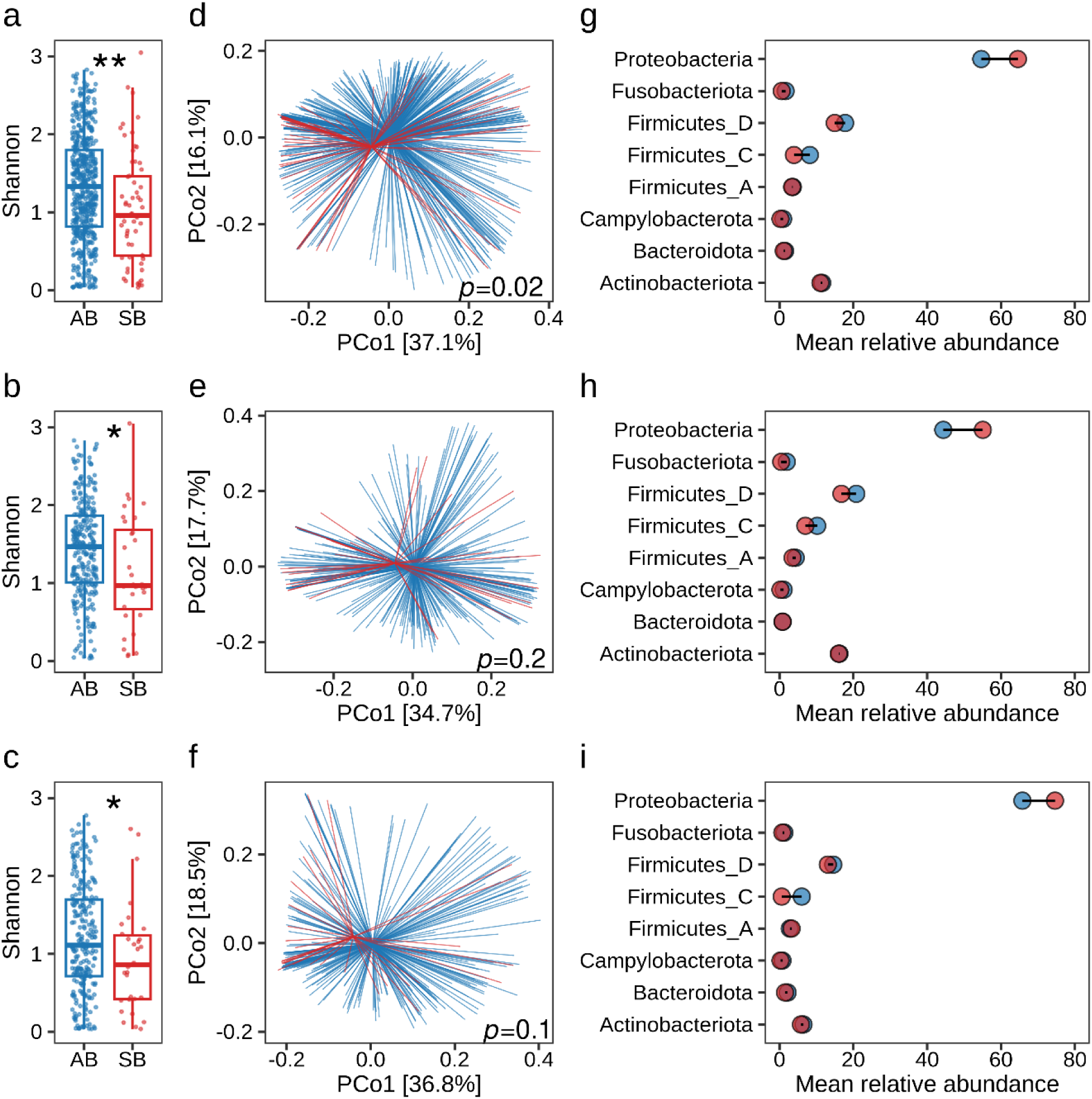
Differences in microbiota community structure and composition between asymptomatic (AB) and symptomatic (SB) bacteriuria samples. (a-c) Alpha diversity measured by the Shannon index of AB (blue) and SB (red) samples across combined (a), catheter (b) and urine (c). Each data point represents an individual sample. Statistical analysis was performed using Wilcoxon rank-sum test and significance is indicated by, ** *P* < 0.01; * *P* < 0.05 between groups. (d-f). Principal coordinates analyses (PCoA) of beta-diversity between groups based on weighted unifrac distance matrices are shown across combined (d), catheter (e) and urine (f). Each group is shown in a different colour (AB: blue, SB: red) with centroid and each line represents an individual sample. Statistical significance was determined by permutational ANOVA (PERMANOVA) with 999 permutations between groups and pairwise p-values are indicated inside of each plot. (g-i) Overview of taxonomic composition in AB and SB groups across combined (g), catheter (h) and urine (i). The points (AB: blue, SB: red) and solid line (black) depicting mean relative abundances in percentages and their differences, respectively, for the phyla as indicated in the y-axis.

Taxonomic analysis was conducted to identify bacterial groups driving the differences between AB and SB samples. At the phylum level, the greatest differences in the mean relative abundances were observed for Proteobacteria, followed by Firmicutes_C and Fusobacteriota (Fig. 3g-I and Supplementary Fig. S4). The mean relative abundance of Proteobacteria was higher in SB (64.5%) compared to AB (54.6%). Conversely, Firmicutes_C (AB: 8.2%, SB: 3.9%) and Fusobacteriota (AB: 1.7%, SB: 0.6%) displayed lower abundances in SB compared to AB. This pattern mirrored the differences observed in the catheter and urine subsets, though with some variations. Notably, Proteobacteria abundance was higher in urine (AB: 65.8%, SB: 74.6%) compared to the catheter (AB: 44.4%, SB: 55.1%). Conversely, Actinobacteriota and Firmicutes displayed higher abundances in the catheter compared to urine. Hence, the phyla level analysis revealed that Proteobacteria and Firmicutes_C were mostly associated with SB and AB respectively. At the family level, Enterobacteriaceae_A, Pseudomonadaceae and Actinomycetaceae were the three most abundant families observed across the three datasets. Genus-level analysis further revealed distinct profiles between AB and SB. Notably, SB samples harboured a higher proportion of *Achromobacter, Actinotignum, Escherichia*_710834, *Massilia, Proteus, Staphylococcus,* and *Stenotrophomonas*_A. In contrast, AB samples showed higher abundances of *Enterococcus*_B, *Fusobacterium*_C, *Serratia*_D, *Streptococcus*, and *Veillonella*_A.

**Fig. 3.**
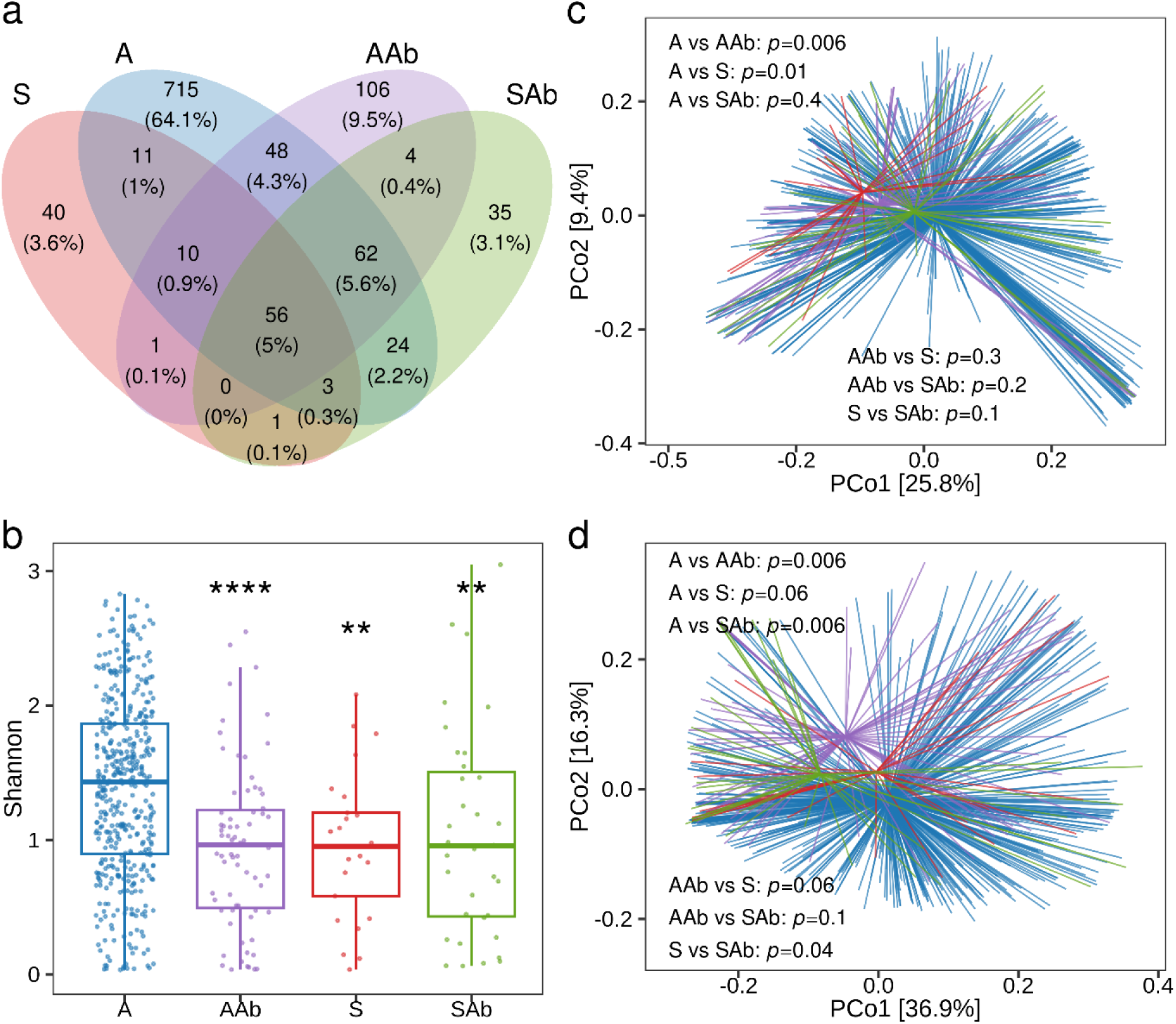
Differences in microbiota community structure among asymptomatic and symptomatic bacteriuria groups with and without antibiotics use. (a) Venn diagram showing the unique and shared ASVs among four groups, asymptomatic samples without antibiotics use (A), asymptomatic samples with antibiotics use (AAb), symptomatic samples without antibiotics (S), and symptomatic samples with antibiotics (SAb). (b) Alpha diversity measured using the Shannon index among the four groups are shown in boxplots. Each data point represents an individual sample. Statistical analysis was performed using Wilcoxon rank-sum test and significance are indicated by, **** *P* < 0.0001; ** *P* < 0.01 compared to group A (c-d). Principal coordinates analyses (PCoA) of beta-diversity between groups based on unweighted (c) and weighted (d) unifrac distance matrices are shown. Each group is shown in a different colour with centroid and each line represents an individual sample. Statistical significance was determined by permutational ANOVA (PERMANOVA) with 999 permutations between groups and pairwise p-values are indicated inside of each plot.

These findings highlight the value of analysing both urine and catheter samples for a more comprehensive understanding of microbial community composition, particularly in identifying UTI-related signatures. While urine alone may reveal some differentiation, combining both sample types provides a more nuanced picture.

### Symptomatic bacteriuria and use of antibiotics lead to alterations in microbial community composition

About one-fifth (∼19%) of the samples (14% of AB and 61% of SB) were collected while participants were taking antibiotics. This use may not always have been for UTI but for other secondary infections. Since antibiotics disrupt the microbiota, we aimed to understand the true differences in microbial community composition between asymptomatic and symptomatic individuals, unaffected by antibiotic influence. We divided the samples based on antibiotic use and UTI symptoms: Asymptomatic without antibiotics (A), Asymptomatic with antibiotics (AAb), Symptomatic without antibiotics (S) and Symptomatic with antibiotics (SAb). These four groups shared 5% (n = 56) of the ASVs while 64.1% (n = 715), 9.5% (n = 106), 3.6% (n = 40) and 3.1% (n = 35) of ASVs were unique to A, AAb, S and SAb, respectively (Fig. 3a). This suggests distinct microbial compositions for each group.

Alpha diversity analysis revealed that both antibiotic use and symptomatic bacteriuria lead to a significant decrease in diversity (Fig. 3b). A significant decrease in alpha diversity was observed in AAb (*P* < 0.0001), S (*P* < 0.01) and SAb (*P* < 0.01) compared to the A group. No significant difference was observed between S and SAb. The community compositions among four groups were also evaluated by unweighted (qualitative) and weighted (quantitative) unifrac beta diversity metrices (Fig. 3c-d). The unweighted measure of beta diversity metrics further demonstrated significant differences between A and S. The unweighted UniFrac showed a significant separation between A vs. AAb (*P* = 0.006) and A vs. S (*P* = 0.01). The weighted UniFrac showed a significant pairwise separation between the antibiotic treated group compared to the A and S.

Analysis of the predominant taxa revealed higher relative abundances of Proteobacteria in antibiotic-treated groups (AAb 64.3% and SAb 71.7%) compared to untreated groups (A 53.1% and S 57.6%) (Fig. 4). This increase can be attributed to the Pseudomonadaceae family, with a mean relative abundance of 22.9% and 22.8 % in AAb and SAb, respectively. Additionally, the S group displayed higher proportions of Firmicutes_D (∼25%) compared to the other sample groups. At the family level, Enterobacteriaceae_A dominated the S group, with the highest mean relative abundance (49.3%). Other notable families in S included Staphylococcaceae (17.9%), Actinomycetaceae (8.9%), Xanthomonadaceae_616009 (3.1%) and Mycobacteriaceae (2.3%). Genus-level analysis revealed enrichment of *Acinetobacter, Actinotignum, Corynebacterium, Escherichia*_710834, *Morganella, Proteus, Staphylococcu*s, and *Stenotrophomonas* in S compared to other sample groups. Notably, four genera: *Escherichia*_710834 (30.3%), *Staphylococcus* (17.9%), *Proteus* (10.6%), and *Actinotignum* (8.9%) constituted over two-thirds of the S group bacterial composition.

**Fig. 4.**
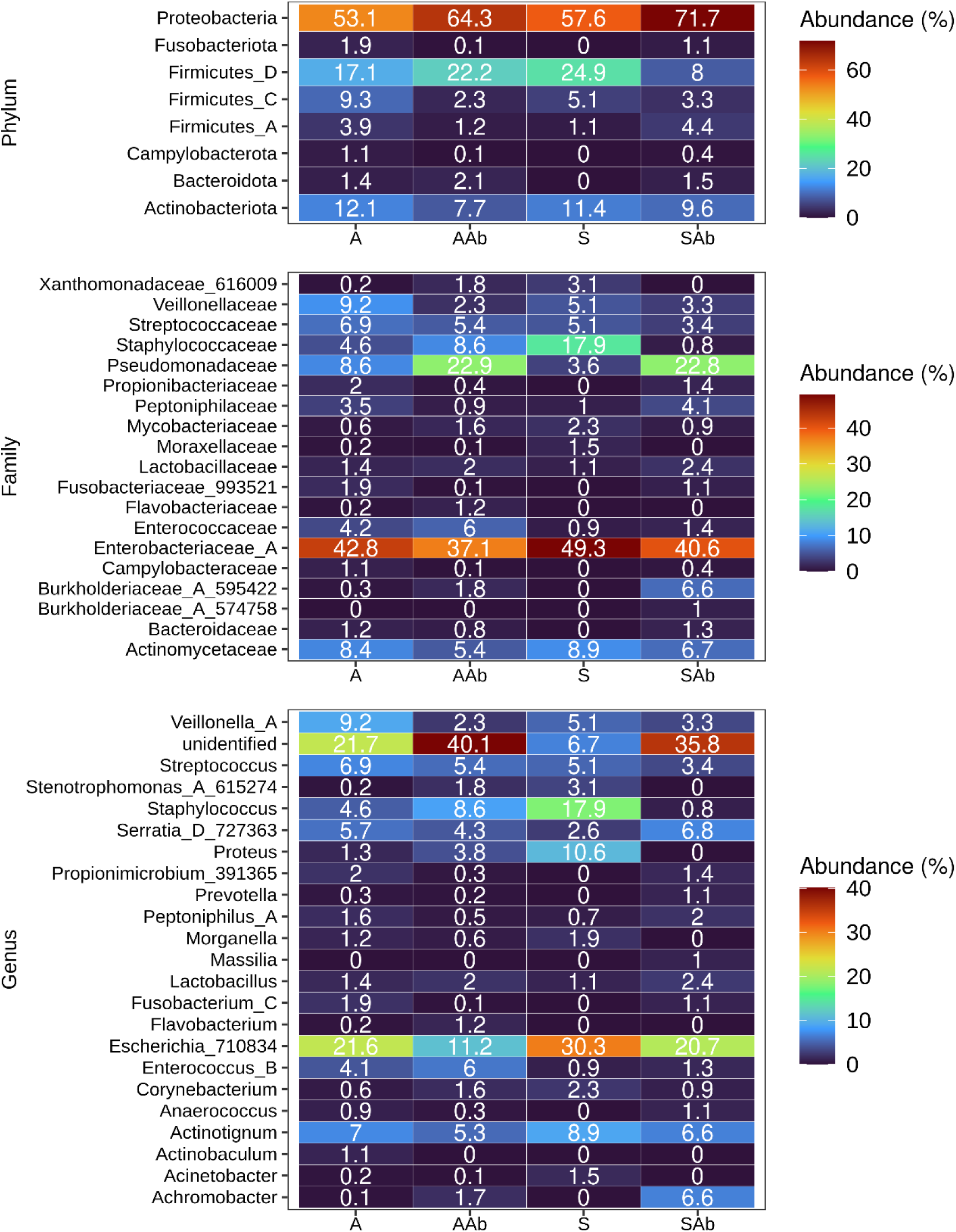
Overview of taxonomic composition in asymptomatic and symptomatic bacteriuria with and without antibiotics treated groups from combined dataset. The colour heatmaps depicting mean relative abundances in percentages ranging from blue (low abundance) to red (high abundance) by phylum, family and genus. The numbers inside the heatmaps show mean relative abundance of corresponding taxa indicated in y-axis across four different groups, asymptomatic samples without antibiotics use (A), asymptomatic samples with antibiotics use (AAb), symptomatic samples without antibiotics (S) and symptomatic samples with antibiotics (SAb). Family and genus are shown if their mean relative abundances in any of the group was more than 1.

These results confirm that community composition between asymptomatic (A) and symptomatic (S) groups differs significantly. The analysis also revealed that the use of antibiotics significantly alters the community composition in asymptomatic samples. These findings also highlight the value of analysing antibiotic treated and untreated samples for a more comprehensive understanding of microbial community composition. Furthermore, this analysis identified potential taxa associated with A and S, highlighting their potential as biomarkers for differentiating these two groups.

### Machine learning can classify symptomatic and asymptomatic bacteriuria with high accuracy

Our study explored the potential of urine and catheter microbial composition as a diagnostic tool for classifying symptomatic and asymptomatic bacteriuria using supervised machine learning. Two feature sets derived from 16S rRNA gene amplicon analysis, ASVs counts and taxonomic abundances (Taxa), were used to train and evaluate prediction models. We aimed to accurately classify both AB and SB patients. Clinically, it is important to determine the timeframe over which patients with AB can retain their existing instilled catheters, while those with SB, or at-risk microbiological profiles, may need catheter replacement to minimise advanced UTI risks. Therefore, we applied the AUROC metric, which evaluates the trade-off between sensitivity and specificity across all possible thresholds, allowing for comprehensive comparisons of classifier performance on various datasets. Recognising the imbalanced nature of our dataset (more AB cases, fewer SB), we additionally provide AUPRC as a complementary measure. AUPRC focused on the trade-off between precision and recall, considering a baseline equivalent to the proportion of minority class (SB) within the entire sample. Both cross-validation and held-out set results were reported since including both results demonstrate the robustness of the model’s performance as the former estimates the stability and generalisability of a model by repeatedly training and testing on different subsets of data, while the later one provides an independent evaluation of the model (Fig. 5a). Additionally, evaluation of both is useful to check any overfitting and underfitting performance of the trained model. Here, the majority of datasets showed a similar level of performance during cross-validation and held-out evaluation (Fig. 5b). The mean AUROC differences between the cross-validation and held-out sets were not statistically different in the majority of datasets. This indicates that the model did not show any overfitting or underfitting issues, particularly in ASVs and without antibiotic datasets.

**Fig. 5.**
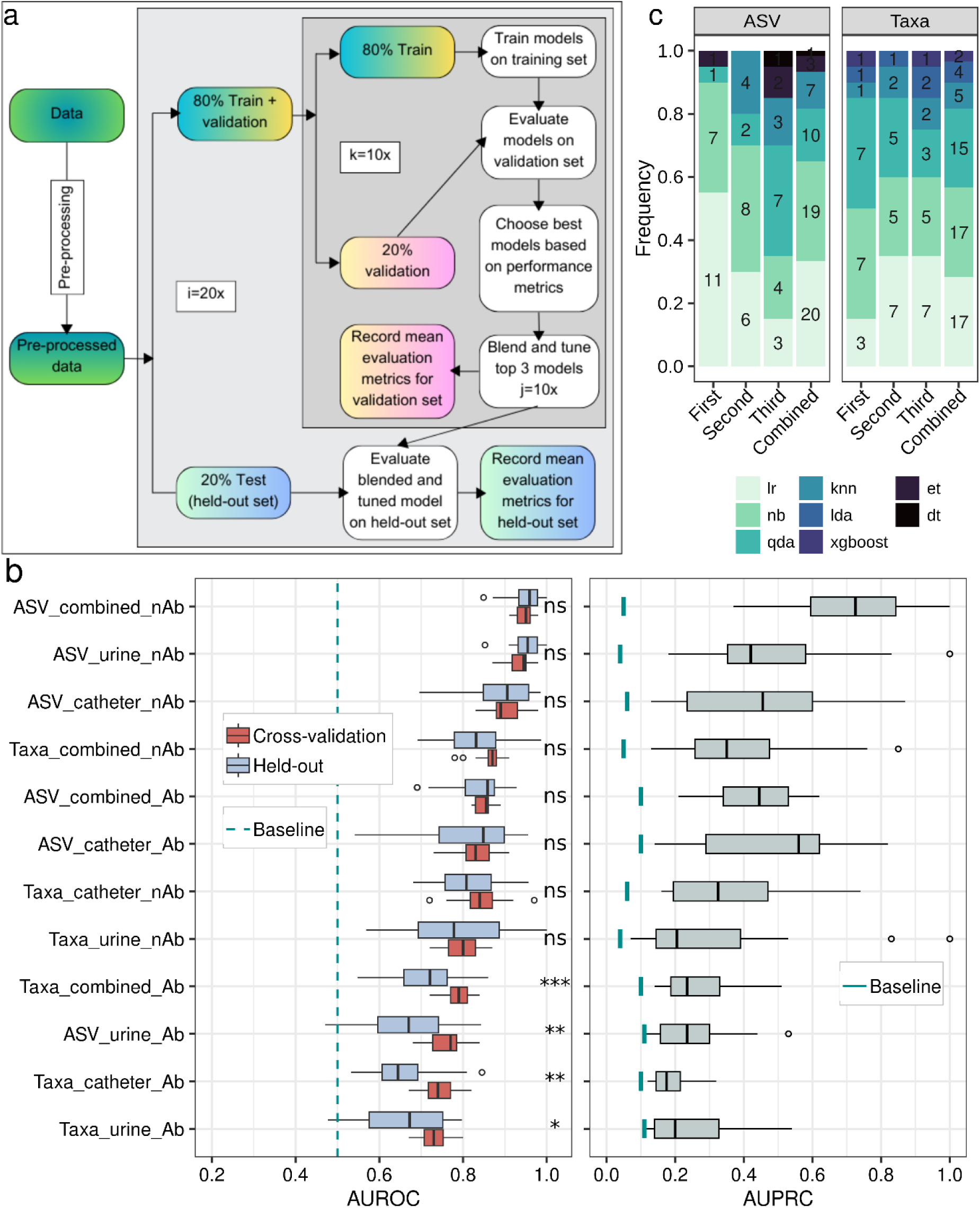
Workflow and predictive performance of machine learning models based on microbiota composition. (a) Workflow for supervised machine learning. The pre-processed data were subjected to stratified (proportional class distribution) split to create 80% training and 20% held-out sets (repeated 20 times). A 10-fold cross-validation was performed on the training data to select the best models. Top three models based on accuracy were blended and tuned (repeated 10 times). The blended and tuned model performance was evaluated on both cross-validation and held-out sets. (b) The boxplots show performance of ML models using AUROC on cross-validation and held-out testing set (left panel) and AUPRC on held-out testing set (right panel) across different datasets. The datasets were arranged in descending order from top to bottom based on the mean AUROC values. The median depicted as centre line in the box, edges depict inter-quartiles, and whiskers as distribution of the data (1.5 times of the quartiles). Outliers are shown as points. The random chances of AUROC depicted by a vertical dashed ‘dark-cyan’ line at 0.5. The baseline chances of AUPRC depicted as vertical solid ‘dark-cyan’ line underneath of the boxplots for each dataset. The baseline performance for AUPRC was calculated as the fraction of the samples in the minority class (SB) over the total number of samples in the test set. Statistical analysis was performed using Wilcoxon rank-sum test and significance is indicated by, *** *P* < 0.001; ** *P* < 0.01; * *P* < 0.05; ns: not significant between cross-validation and held-out set. (c) Frequency of the top three blended models across ASV and Taxa (combined and without antibiotics) datasets.

Model performance was evaluated across three variables: antibiotic use (samples with vs. without), feature type (ASVs vs Taxa) and sample type (catheter, urine or combined). Firstly, we assessed model performance on samples with and without antibiotic treatment. This aimed to account for the significant effect antibiotics have on bacterial diversity. Importantly, diagnosing untreated samples (person) seeking medical advice is more clinically relevant. However, we also evaluated samples with antibiotic use, considering that many SCI patients receive antibiotics, necessitating accurate diagnosis of both antibiotic-associated asymptomatic and symptomatic bacteriuria in these cases. Cross-validation results showed that excluding antibiotic-treated samples improved model accuracy. Mean AUROC scores with ASV features ranged from 0.91-0.98 (IQR 0.93-0.96) without antibiotics to 0.82-0.89 (IQR 0.83-0.86) with antibiotics (Fig. 5b and Supplementary Tables S3). Held-out set evaluation confirmed this trend, with a mean AUROC of 0.85-1 (IQR 0.93-0.98) and 0.69-0.93 (IQR 0.81-0.88) in untreated and treated samples, respectively (Fig. 5b and Supplementary Tables S4). The ASV feature on combined and without antibiotic dataset showed the highest AUPRC with a mean of 0.37-0.1 (IQR 0.6-0.84) compared to any other dataset with a baseline AUPRC value of 0.05. These findings suggest that excluding antibiotic-treated samples improves overall model performance for combined (catheter and urine) datasets.

Next, we compared model performance between ASV and taxa features and found that ASV yielded higher AUROC scores, reaching 0.91-0.98 (IQR 0.93-0.96) compared to 0.78-0.91 (IQR 0.86-0.88) for taxa when trained on untreated combined datasets (Fig. 5b and Supplementary Tables S3). Held-out set evaluation also showed this trend, with a mean AUROC of 0.85-1 (IQR 0.93-0.98) and 0.69-0.99 (IQR 0.78-0.88) for ASV and taxa features, respectively (Fig. 5b and Supplementary Tables S4). The same trend was also observed in datasets with antibiotics, which showed better performance with ASV during cross-validation with mean AUROC of 0.82-0.89 (IQR 0.83-0.86) compared to taxa (AUROC 0.72-0.84, IQR 0.77-0.81) datasets (Fig. 5b and Supplementary Tables S3). Held-out evaluation showed a mean AUROC of 0.69-0.93 (IQR 0.81-0.88) and 0.55-0.86 (IQR 0.66-0.76) for ASV and taxa feature, respectively (Fig. 5b and Supplementary Tables S4). The ASV feature also showed a higher AUPRC score compared to taxa on both with and without antibiotic dataset. Hence, the results show in general a better performance of the models trained with ASV feature compared to the taxa feature across all datasets.

Finally, we compared model performance on combined datasets versus catheter and urine-only datasets, considering the differences in diversity observed between these groups. ASV features and exclusion of antibiotic-treated samples led to the best performance in combined datasets (mean AUROC 0.91-0.98, IQR 0.93-0.96) compared to catheter (AUROC 0.83-0.98, IQR 0.88-0.93) or urine-only datasets (AUROC 0.87-0.98, IQR 0.92-0.95) during cross-validation (Fig. 5b and Supplementary Tables S3). Held-out evaluation also confirmed this trend, with ASV features achieving a mean AUROC of 0.85-1 (IQR 0.93-0.98) in the combined dataset, compared to 0.7-0.99 (IQR 0.85-0.96) for catheter and 0.85-1 (IQR 0.93-0.98) for urine-only datasets (Fig. 5b and Supplementary Tables S4). The AUPRC score was highest for the combined dataset 0.37 to 1 (IQR 0.6-0.84) followed by urine 0.18 to 1 (IQR 0.35-0.58) and catheter 0.13 to 0.87 (IQR 0.24-0.6) with a baseline 0.05, 0.04 and 0.06 respectively. These findings indicate that using combined datasets significantly improved model performance compared to analysing individual sampling sites.

We blended the top three performing models and evaluated the performance on the blended and tuned model. Among the 16 tested models, we observed that nine models appeared in the top three list when evaluated in ASV and Taxa combined and without antibiotics datasets (Fig. 5c). Logistic regression (lr), Naïve bayes (nb) and Quadratic discriminant analysis (qda) classifier were the highest performing classifier.

### ASVs and Taxa belonging to Proteobacteria phyla showed the highest importance in classification of AB and SB

Given the good predictive performances of the models trained on the ASV feature, we next sought to identify ASVs that were most important in classifying the AB and SB using the feature importance derived from the top performing classifier. We plotted the top 20 ASVs, of which 9 ASVs belonged to the Proteobacteria phyla, which includes members of 8 Enterobacteriaceae_A and 1 Pseudomonadaceae families (Fig. 6a). A member of the *Escherichia*_710834 genus (ASV 1126) had the strongest effect on feature importance followed by a member of the *Staphylococcus* genus (ASV 224). Plotting the relative abundance of these top 20 ASVs revealed significant differences between AB and SB (Fig. 6a). In particular, the median relative abundance of the genus *Escherichia*_710834 (ASV 1126, 1040, 1074) was higher in SB compared to AB. The relative abundance of the genus *Escherichia*_710834 (ASV 1020), Enterobacteriaceae_A (ASV 933), *Enterococcus_B* (ASV 163) and *Staphylococcus* (ASV 224) were significantly different between AB and SB (*P* < 0.05). The ASV 1126 and 205 were unique to SB corresponding to *Escherichia_*710834 and *Staphylococcus* genus, respectively. In contrast, seven ASVs were unique to AB, ASV 292, 412, 456, 696, 936, 1005 and 1019 corresponding to *Streptococcus constellatus, Fastidiosipila sanguinis*, *Veillonella* genus, *Campylobacter ureolyticus* and Enterobacteriaceae_A family, respectively.

**Fig. 6.**
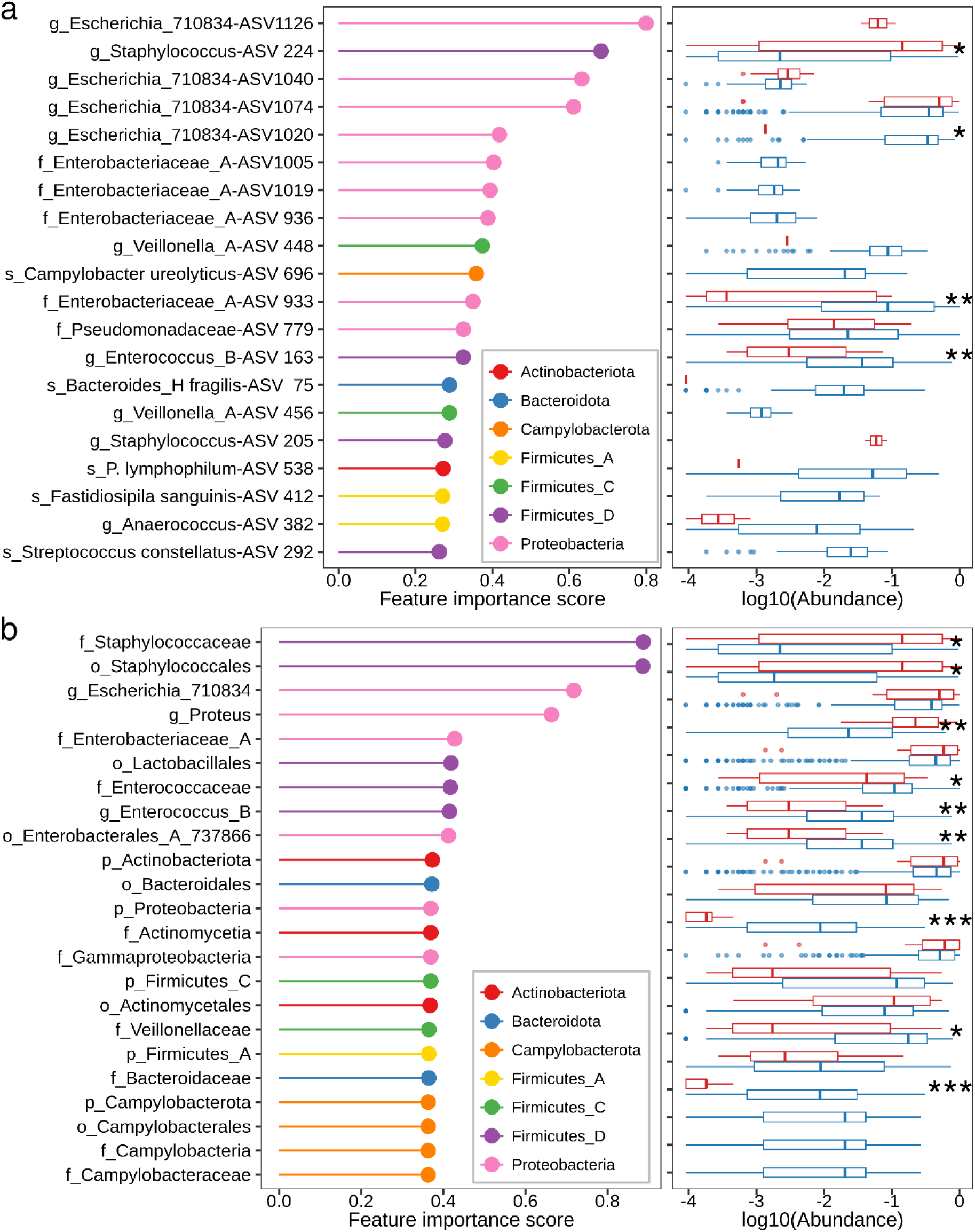
Important ASVs and taxa features contributing to the classification of AB and SB. Feature importance of the top 20 most important ASVs (a) and Taxa (b) derived from the top performing classifier. Colour represents the phyla corresponding to each ASVs and Taxa. The right panel depicts the differences in log10-transformed relative abundance for the top 20 most important ASVs (a) and Taxa (b) between symptomatic and asymptomatic bacteriuria samples. Statistical analysis was performed using the Wilcoxon rank-sum test and significance is indicated as *** *P* < 0.001; ** *P* < 0.01; * *P* < 0.05 between groups.

In addition to the ASV feature, we also sought to identify taxa that were most important in classifying the AB and SB using the feature importance derived from the top performing classifier. Interestingly, many taxa identified were similar to the ASV analysis, and 6 out of the top 20 taxa belonged to the Proteobacteria phyla (Fig. 6b). Among the top 20 taxa, the family Staphylococcaceae had the strongest effect, followed by the three members of the Proteobacteria phyla. At the genus level, *Escherichia*_710834 showed the highest effect, followed by *Proteus* and *Enterococcus*. In the majority of cases, the feature importance score corresponded to the differences in mean abundances of these taxa observed during taxonomic analysis (Fig. 4).

## Discussion

SCI individuals are often critically ill and may require long-term catheter use for urination, which can lead to an increased risk of developing bacteriuria and CAUTI [45]. Frequent catheter changes are unpleasant for patients and can be painful. Therefore, it is crucial to monitor these patients closely to determine the optimal catheter change schedule, balancing infection prevention and minimising unnecessary procedures. A key clinical challenge lies in differentiating asymptomatic and symptomatic bacteriuria due to the reported presence of UTI-causing pathogens in both states [7, 46, 47]. Additionally, the likelihood of colonisation and biofilm formation progressing to clinical infection is often related to patient-specific immunological background, the types of catheter biomaterial, microbiota present as well as environmental and medication factors [48–50].

Our study also corroborates the previous findings that many pathogens overlap between asymptomatic and symptomatic bacteriuria states. Moreover, previous studies have also shown that infections disrupt the urinary and catheter microbiome, causing an imbalance in the normal bacterial community and allowing pathogens to dominate [14, 51, 52]. Similar to those studies, our results indicate that symptomatic bacteriuria and antibiotic use are associated with distinct microbial communities compared to healthy states. This is reflected in our diversity analysis, which revealed lower alpha diversity (species richness) in symptomatic samples and differences in beta diversity (community composition) between asymptomatic and symptomatic samples. We also observed increased abundance of UTI-associated pathogens in these samples. These findings align with our previous pilot study, suggesting that community composition changes in response to disruptions, such as antibiotic treatment or pathogen colonisation, which can lead to CAUTI [14]. Here, we demonstrated that SB alters the microbial community structure in patients with SCI. These changes are associated with an increase in abundance of members of the *Escherichia* sp., *Staphylococcus* sp., *Proteus* sp., *Actinotignum* sp., *Corynebacterium* sp., and *Morganella* sp. genus. These bacteria are known to be major causes of UTI and also have been previously identified in urine samples [7, 47, 53].

This study investigated machine learning approaches that utilises microbial signatures to classify AB and SB in patients with SCI. We demonstrated the effectiveness of this approach across various sample types, including samples with and without antibiotic treatment and those obtained from catheters and urine. The model achieved the highest performance with samples that had not received antibiotic treatment. In these samples the model could predict AB and SB with over 90% accuracy and at 7-20 times greater precision compared to the baseline precision. While diagnosing untreated samples holds greater clinical relevance, a significant portion of the SCI population require antibiotics. These antibiotics may not always target UTI-causing pathogens but address secondary infections or complications. Even including samples that received antibiotics, the model maintained over 80% accuracy and achieved 2-6 times greater precision compared to the baseline in predicting AB and SB. Therefore, our study explored the suitability of the model in both scenarios, demonstrating its ability to classify AB and SB with high accuracy regardless of antibiotic treatment.

Our highest performing model utilised both catheter and urine samples for prediction. However, we observed that urine samples alone yielded better performance compared to catheter samples in identifying AB and SB. This is advantageous because fresh-catch urine samples are easier to obtain and evaluate for screening purposes. The improved performance of urine-based models might be attributed to the higher abundance of Proteobacteria phyla identified through taxonomic analysis. Our data showed approximately 20% increase in Proteobacteria in urine samples with SB compared to catheter. Future investigations are needed to definitively determine why urine is a more informative sample type than the catheter biofilm content. Potential explanations include the frequent route of exposure for pathogens in the urinary tract and the bladder serving as a more suitable niche for UTI-causing bacteria compared to the biofilms of a catheter. Additionally, UTI pathogens may be more motile or dispersive, and bacteria may adhere to and colonise at different rate in catheters compared to the urinary tract and bladder. In support of these statements, a previous study has shown increased association of the members of Proteobacteria phyla particularly *E. coli* and *Klebsiella* sp. in urine samples compared to catheter biofilm contents in SCI patients [54].

Our study demonstrated that ASVs offer greater advantages over taxa features for machine learning tasks in predicting AB and SB. The underlying advantages of ASV features over taxa might be attributed to the higher resolution, improved accuracy and strain level insights provided by ASVs. Compared to taxa, ASVs exhibit subtle sequence variations. Often, multiple ASVs can map to a single taxon, providing a more precise picture of microbial communities. Furthermore, understanding strain-level variation within a species is crucial, as these closely related strains might have distinct functional roles in the microbiome. ASV based analysis allowed us to identify and differentiate between such strains. Interestingly, our results revealed that specific groups of ASVs belonging to the same taxon were enriched in either SB or AB. This suggests that particular pathogenic strains might predominate in each state. Future studies with strain-level resolution in samples from AB and SB are necessary to confirm this. In addition, machine learning algorithms perform best with informative features. Since ASVs capture finer genetic variations, they offer a richer signal for the model to learn from. This potentially leads to more accurate predictions compared to broader taxonomic classifications. While our analysis showed improved performance using ASVs, it’s important to acknowledge the success of the taxa-based approach as well. Taxa-based models achieved an average accuracy exceeding 80% and a 3-17 fold increase in precision compared to the baseline.

In this study, we employed PyCaret to build ensemble models by selecting the top three performing machine learning classifiers out of the sixteen evaluated. Since individual classifiers often excel at predicting specific classes but struggle with others, combining them improves overall prediction accuracy for both classes. This approach, utilising a soft voting system, significantly enhances model performance compared to single classifiers. While ensemble models are not a new concept, their application in disease diagnosis remains limited. Most existing literature focuses on individual models for disease classification [20, 55]. However, our study, along with others employing ensemble models, demonstrates the growing applicability and effectiveness of this approach [18, 56]. This success suggests that similar ensemble strategies could be implemented to achieve high-accuracy classification in other disease states.

Our study has several limitations. First, potential under- and over-reporting of asymptomatic and symptomatic events may have occurred. Self-reported symptomatic events without confirmatory pathology could lead to overestimation, while chronic UTI patients may tolerate or ignore symptoms, causing underestimation. Second, catheter samples were collected only during routine changes due to the invasive nature of the procedure. More frequent sampling would have provided valuable insights into microbial dynamics and potentially improved model performance. Since our model performed better with urine samples compared to catheter, future studies could collect and analyse urine samples for prediction which, unlike catheters, is easier to obtain and allows for more frequent sampling. Third, our study did not include data on patient dietary habits and other potential immunological and environmental inputs in the analysis. Finally, the training data for our machine learning model included a relatively small cohort of symptomatic samples. We addressed class imbalance using techniques like stratified splitting and SMOTE and reported metrics that account for this. While our model showed promise within the recruited cohort, further validation with a larger, external cohort is crucial. This will help identify potential biases that might affect its performance in a clinical setting. There is also a need to determine whether including other potential biomarkers and indicators of immunological resilience and environmental risks of potential infection could improve the accuracy of the current model. In addition, external validation is essential to confirm the model’s effectiveness. However, acquiring such data can be challenging due to limited metadata associated with existing sequence and the need for the exact variable regions of the 16S rRNA gene that was used in this study.

## Conclusion

Our findings reveal several unique characteristics of symptomatic bacteriuria in SCI patients, including lower microbial diversity, compositional changes, and enrichment of UTI associated pathogens. This study represents the first comprehensive microbiome profiling of both catheter and urine samples from SCI patients and we utilised this data to develop a machine learning model for UTI prediction. While future inclusion of more samples could improve the model’s class balance, the current version demonstrates high accuracy and holds promise for real-world healthcare implementation. This could significantly improve patient quality of life and guide treatment decisions. We demonstrated that 16S rRNA amplicon sequencing data could be used to predict asymptomatic and symptomatic bacteriuria with high accuracy. These results have significant implications for establishing an early warning system for potential UTI in SCI patients. The benefits of our model are threefold. First, it can predict potential UTI, informing decisions about catheter changes to prevent potential infections. Second, for patients predicted to have asymptomatic bacteriuria, the model can recommend keeping the catheter, reducing unnecessary procedures and costs. Third, it can prevent unnecessary antibiotic use, thereby curbing the rise of multidrug-resistant bacteria.

Overall, this study evaluated the diagnostic potential of machine learning models for future implementation in treatment decisions and intervention strategies to better protect this high-risk patient population. Looking forward, we aim to implement our model into healthcare settings to classify asymptomatic and symptomatic bacteriuria in SCI patients. This has the potential to improve patient quality of life, reduce mortality rates, curb the spread of drug-resistant bacteria and generate significant cost savings for hospitals.

## Abbreviations

SCI: Spinal cord injury.
UTI: Urinary tract infection.
CAUTI: Catheter associated urinary tract infections.
DNA: Deoxyribonucleic acid.
rRNA: Ribosomal Ribonucleic acid.
PCR: Polymerase chain reaction.
ASVs: Amplicon sequence variants.
PCoA: Principal coordinate analysis.
PERMANOVA: Permutational multivariate analysis of variance.
QIIME: Quantitative insights into microbial ecology.
IQR: Interquartile range.
SMOTE: Synthetic minority over-sampling technique.
AUROC: Area under the receiver operating characteristic curve.
AUPRC: Area under the precision-recall curve.

## Supplementary Information

**Additional file 1 (.pdf):**

**Supplementary Fig. S1.** Treemap showing the fraction of ASVs assigned to Family. The number and percentages of ASVs belong to each family are denoted inside of each box. The colors depict corresponding phyla.

**Supplementary Fig. S2.** Beta diversity (unweighted unifrac) analysis in participants with at least one symptomatic bacteriuria event. Principal coordinates analyses (PCoA) of beta-diversity between asymptomatic (AB, blue) and symptomatic (SB, red) bacteriuria group based on unweighted unifrac distance matrices. Each dot represents an individual sample.

**Supplementary Fig. S3.** Beta diversity analysis (weighted unifrac) in participants with at least one symptomatic bacteriuria event. Principal coordinates analyses (PCoA) of beta-diversity between asymptomatic (AB, blue) and symptomatic (SB, red) bacteriuria group based on weighted unifrac distance matrices. Each dot represents an individual sample.

**Supplementary Fig. S4**. Overview of taxonomic composition in asymptomatic and symptomatic bacteriuria groups across combined, catheter and urine dataset. The colour heatmaps depicting mean relative abundances in percentages ranging from blue (low abundance) to red (high abundance) grouped by phylum, family, and genus. The numbers inside heatmaps show mean relative abundance of corresponding taxa indicated in y-axis across asymptomatic bacteriuria (AB) and symptomatic (SB) groups. Genus are shown if their mean relative abundances in any of the group was more than 1.

**Supplementary Table S1**. Participants information and samples analysed.

**Supplementary Table S2.** Machine learning models.

**Supplementary Table S3.** Performance of machine learning models during cross-validation.

**Supplementary Table S4.** Performance of machine learning models on held-out set.

## Declarations

### Ethics approval and consent to participate

The samples for this project were obtained as part of a multi-centre observational longitudinal study 3PU (ACTRN12622000613707). The ethics approval for all study protocols were obtained from the Northern Sydney Local Health District Human Research Ethics Committee (HREC) 2021/ETH00329 as well as Site Specific Assessment (SSA) 2021/STE00542 by Royal North Shore Hospital governance and 2021/STE00541 by Prince of Wales Hospital governance. Participants Adults, age 18 years and older, with stable SCI and stable neurogenic bladder management technique for at least 4 weeks before start of the study. Participants agreed to have their urinary system microbial community and the microbiome DNA from the catheter and urine to be extracted and stored long-term. Written informed consent was obtained from all participants and samples were de-identified by the assigning of an arbitrary subject number.

### Consent for publication

Not applicable

### Author’s contributions

DM, SAR, IGD and BBL developed the concept and designed the study. PN, JT, PC, GW, JP and BBL co-ordinate the recruitment of all participants, collection of the samples, and fortnightly telephone conversation. JT, PN, GEV, MMH and DL processed samples. JT extracted DNA from all samples and performed 16S rRNA gene sequencing. MMH processed the 16S rRNA gene sequencing data, performed bioinformatics, statistical, and machine learning analyses. MMH, GEV, PN wrote the manuscript with input from all authors. DM, SAR, IGD and BBL supervised the project. All authors provided valuable feedback to improve the final manuscript.

## Acknowledgement

We thank all the participants and care team who sincerely provided us the samples, especially clinical nurses Ruth Hamilton, Ruyi Yao and Jennifer Greenway. We also like to thank staff at Ferguson lodge and staff of Fairfield west clinic and Dr. Zahra Rassoly Obayd for their help with participant recruitment and sample collection. We would like to thank Dr. Obaydullah Marial for his advice on design of our participant data collection forms and questionaries. We also thankful to UTS high performance computing facility iHPC for providing us the platform for data analysis.

## References

1. Skelton-Dudley F, Doan J, Suda K, Holmes SA, Evans C, Trautner B. Spinal cord injury creates unique challenges in diagnosis and management of catheter-associated urinary tract infection. Topics in spinal cord injury rehabilitation. 2019;25(4):331–9.

2. Dalen DM, Zvonar RK, Jessamine PG. An evaluation of the management of asymptomatic catheter-associated bacteriuria and candiduria at The Ottawa Hospital. Canadian Journal of Infectious Diseases and Medical Microbiology. 2005;16:166–70.

3. Nicolle LE. Catheter associated urinary tract infections. Antimicrobial resistance and infection control. 2014;3:1–8.

4. Nicolle LE. Asymptomatic bacteriuria: when to screen and when to treat. Infectious Disease Clinics. 2003;17(2):367–94.

5. Trautner BW, Cope M, Cevallos ME, Cadle RM, Darouiche RO, Musher DM. Inappropriate treatment of catheter-associated asymptomatic bacteriuria in a tertiary care hospital. Clinical Infectious Diseases. 2009;48(9):1182–8.

6. Rubi H, Mudey G, Kunjalwar R. Catheter-associated urinary tract infection (CAUTI). Cureus. 2022;14(10).

7. Fouts DE, Pieper R, Szpakowski S, Pohl H, Knoblach S, Suh M-J, et al. Integrated next-generation sequencing of 16S rDNA and metaproteomics differentiate the healthy urine microbiome from asymptomatic bacteriuria in neuropathic bladder associated with spinal cord injury. Journal of translational medicine. 2012;10:1–17.

8. Werneburg GT. Catheter-associated urinary tract infections: current challenges and future prospects. Research and reports in urology. 2022:109–33.

9. Hollenbeak CS, Schilling AL. The attributable cost of catheter-associated urinary tract infections in the United States: A systematic review. American journal of infection control. 2018;46(7):751–7.

10. McCleskey SG, Shek L, Grein J, Gotanda H, Anderson L, Shekelle PG, et al. Economic evaluation of quality improvement interventions to prevent catheter-associated urinary tract infections in the hospital setting: a systematic review. BMJ quality & safety. 2022;31(4):308–21.

11. Podkovik S, Toor H, Gattupalli M, Kashyap S, Brazdzionis J, Patchana T, et al. Prevalence of catheter-associated urinary tract infections in neurosurgical intensive care patients–the overdiagnosis of urinary tract infections. Cureus. 2019;11(8).

12. Flores-Mireles AL, Walker JN, Caparon M, Hultgren SJ. Urinary tract infections: epidemiology, mechanisms of infection and treatment options. Nature reviews microbiology. 2015;13(5):269–84.

13. Albu S, Voidazan S, Bilca D, Badiu M, Truta A, Ciorea M, et al. Bacteriuria and asymptomatic infection in chronic patients with indwelling urinary catheter: The incidence of ESBL bacteria. Medicine. 2018;97(33):e11796.

14. Bossa L, Kline K, McDougald D, Lee BB, Rice SA. Urinary catheter-associated microbiota change in accordance with treatment and infection status. PLoS One. 2017;12(6):e0177633.

15. Foxman B, Wu J, Farrer EC, Goldberg DE, Younger JG, Xi C. Early development of bacterial community diversity in emergently placed urinary catheters. BMC research notes. 2012;5:1–7.

16. Nye TM, Zou Z, Obernuefemann CL, Pinkner JS, Lowry E, Kleinschmidt K, et al. Microbial co-occurrences on catheters from long-term catheterized patients. Nature communications. 2024;15(1):61.

17. Alowais SA, Alghamdi SS, Alsuhebany N, Alqahtani T, Alshaya AI, Almohareb SN, et al. Revolutionizing healthcare: the role of artificial intelligence in clinical practice. BMC medical education. 2023;23(1):689.

18. Chen H-C, Liu Y-W, Chang K-C, Wu Y-W, Chen Y-M, Chao Y-K, et al. Gut butyrate-producers confer post-infarction cardiac protection. Nature communications. 2023;14(1):7249.

19. Barron MR, Sovacool KL, Abernathy-Close L, Vendrov KC, Standke AK, Bergin IL, et al. Intestinal inflammation reversibly alters the microbiota to drive susceptibility to *Clostridioides difficile* colonization in a mouse model of colitis. Mbio. 2022;13(4):e01904–22.

20. Caussy C, Tripathi A, Humphrey G, Bassirian S, Singh S, Faulkner C, et al. A gut microbiome signature for cirrhosis due to nonalcoholic fatty liver disease. Nature communications. 2019;10(1):1406.

21. Forbes JD, Chen C-y, Knox NC, Marrie R-A, El-Gabalawy H, de Kievit T, et al. A comparative study of the gut microbiota in immune-mediated inflammatory diseases—does a common dysbiosis exist? Microbiome. 2018;6:1–15.

22. Ozkan IA, Koklu M, Sert IU. Diagnosis of urinary tract infection based on artificial intelligence methods. Computer methods and programs in biomedicine. 2018;166:51–9.

23. Burton RJ, Albur M, Eberl M, Cuff SM. Using artificial intelligence to reduce diagnostic workload without compromising detection of urinary tract infections. BMC medical informatics and decision making. 2019;19:1–11.

24. Gadalla AA, Friberg IM, Kift-Morgan A, Zhang J, Eberl M, Topley N, et al. Identification of clinical and urine biomarkers for uncomplicated urinary tract infection using machine learning algorithms. Scientific reports. 2019;9(1):19694.

25. Jeng S-L, Huang Z-J, Yang D-C, Teng C-H, Wang M-C. Machine learning to predict the development of recurrent urinary tract infection related to single uropathogen, *Escherichia coli*. Scientific reports. 2022;12(1):17216.

26. Toh S-L, Lee BB, Ryan S, Simpson JM, Clezy K, Bossa L, et al. Probiotics [LGG-BB12 or RC14-GR1] versus placebo as prophylaxis for urinary tract infection in persons with spinal cord injury [ProSCIUTTU]: a randomised controlled trial. Spinal Cord. 2019;57(7):550–61.

27. Lee BB, Toh S-L, Ryan S, Simpson JM, Clezy K, Bossa L, et al. Probiotics [LGG-BB12 or RC14-GR1] versus placebo as prophylaxis for urinary tract infection in persons with spinal cord injury [ProSCIUTTU]: a study protocol for a randomised controlled trial. BMC urology. 2016;16:1–8.

28. Lee B, Haran M, Hunt L, Simpson J, Marial O, Rutkowski S, et al. Spinal-injured neuropathic bladder antisepsis (SINBA) trial. Spinal cord. 2007;45(8):542–50.

29. Bolyen E, Rideout JR, Dillon MR, Bokulich NA, Abnet CC, Al-Ghalith GA, et al. Reproducible, interactive, scalable and extensible microbiome data science using QIIME 2. Nature biotechnology. 2019;37(8):852–7.

30. Martin M. Cutadapt removes adapter sequences from high-throughput sequencing reads. EMBnet journal. 2011;17(1):10–2.

31. Callahan BJ, McMurdie PJ, Rosen MJ, Han AW, Johnson AJA, Holmes SP. DADA2: High-resolution sample inference from Illumina amplicon data. Nature methods. 2016;13(7):581–3.

32. Robeson MS, O’Rourke DR, Kaehler BD, Ziemski M, Dillon MR, Foster JT, et al. RESCRIPt: Reproducible sequence taxonomy reference database management. PLoS computational biology. 2021;17(11):e1009581.

33. McDonald D, Jiang Y, Balaban M, Cantrell K, Zhu Q, Gonzalez A, et al. Greengenes2 unifies microbial data in a single reference tree. Nature biotechnology. 2023:1–4.

34. Katoh K, Standley DM. MAFFT multiple sequence alignment software version 7: improvements in performance and usability. Molecular biology and evolution. 2013;30(4):772–80.

35. Price MN, Dehal PS, Arkin AP. FastTree 2–approximately maximum-likelihood trees for large alignments. PloS one. 2010;5(3):e9490.

36. McMurdie PJ, Holmes S. phyloseq: an R package for reproducible interactive analysis and graphics of microbiome census data. PloS one. 2013;8(4):e61217.

37. Schloss PD. Rarefaction is currently the best approach to control for uneven sequencing effort in amplicon sequence analyses. Msphere. 2024:e00354–23.

38. Lahti L, Shetty S. Tools for microbiome analysis in R. http://microbiome.github.com/microbiome (2017). Accessed 10.09.2023.

39. Liu C, Cui Y, Li X, Yao M. microeco: an R package for data mining in microbial community ecology. FEMS microbiology ecology. 2021;97(2):fiaa255.

40. Hadley W. ggplot2: Elegant Graphics for Data Analysis. https://ggplot2.tidyverse.org (2016). Accessed 15.02.2023.

41. Topçuoğlu BD, Lapp Z, Sovacool KL, Snitkin E, Wiens J, Schloss PD. mikropml: user-friendly R package for supervised machine learning pipelines. Journal of open source software. 2021;6(61).

42. Ali M. PyCaret: An open source, low-code machine learning library in Python. https://www.pycaret.org (2020). Accessed 20.10.2023.

43. Pedregosa F, Varoquaux G, Gramfort A, Michel V, Thirion B, Grisel O, et al. Scikit-learn: Machine learning in Python. the Journal of machine Learning research. 2011;12:2825–30.

44. Anderson MJ. Permutational multivariate analysis of variance (PERMANOVA). Wiley statsref: statistics reference online. 2014:1–15.

45. Esclarín De Ruz A, García Leoni E, Herruzo Cabrera R. Epidemiology and risk factors for urinary tract infection in patients with spinal cord injury. The Journal of urology. 2000;164(4):1285–9.

46. Nally E, Groah SL, Pérez-Losada M, Caldovic L, Ljungberg I, Chandel NJ, et al. Identification of Burkholderia fungorum in the urine of an individual with spinal cord injury and augmentation cystoplasty using 16S sequencing: copathogen or innocent bystander? Spinal Cord Series and Cases. 2018;4(1):85.

47. Adu-Oppong B, Thänert R, Wallace MA, Burnham C-AD, Dantas G. Substantial overlap between symptomatic and asymptomatic genitourinary microbiota states. Microbiome. 2022;10(1):6.

48. Donlan RM. Biofilms: microbial life on surfaces. Emerging infectious diseases. 2002;8(9):881.

49. Wu S, Zhang B, Liu Y, Suo X, Li H. Influence of surface topography on bacterial adhesion: A review. Biointerphases. 2018;13(6).

50. Zou Z, Potter RF, McCoy WH, Wildenthal JA, Katumba GL, Mucha PJ, et al. *E. coli* catheter-associated urinary tract infections are associated with distinctive virulence and biofilm gene determinants. JCI insight. 2023;8(2).

51. Marshall CW, Kurs-Lasky M, McElheny CL, Bridwell S, Liu H, Shaikh N. Performance of conventional urine culture compared to 16S rRNA gene amplicon sequencing in children with suspected urinary tract infection. Microbiology Spectrum. 2021;9(3):e01861–21.

52. Pallares-Mendez R, Cervantes-Miranda DE, Gonzalez-Colmenero AD, Ochoa-Arvizo MA, Gutierrez-Gonzalez A. A Perspective of the urinary microbiome in lower urinary tract infections—A Review. Current Urology Reports. 2022;23(10):235–44.

53. Kim DS, Lee JW. Urinary tract infection and microbiome. Diagnostics. 2023;13(11):1921.

54. Garcia-Marques FJ, Zakrasek E, Bermudez A, Polasko AL, Liu S, Stoyanova T, et al. Proteomics analysis of urine and catheter-associated biofilms in spinal cord injury patients. American Journal of Clinical and Experimental Urology. 2023;11(3):206.

55. Zuo W, Wang B, Bai X, Luan Y, Fan Y, Michail S, et al. 16S rRNA and metagenomic shotgun sequencing data revealed consistent patterns of gut microbiome signature in pediatric ulcerative colitis. Scientific Reports. 2022;12(1):6421.

56. Chandrasekhar N, Peddakrishna S. Enhancing heart disease prediction accuracy through machine learning techniques and optimization. Processes. 2023;11(4):1210.

