## Supplementary Information for "Prediction of symptomatic and asymptomatic bacteriuria in spinal cord injury patients using machine learning"

Running Title: Machine learning based prediction of bacteriuria.

**Keywords:** Spinal cord injury, Bacteriuria, Urinary tract infections, 16S rRNA, Catheter, Urine, Machine learning, Prediction.

**Content:**

Supplementary Fig. S1-4

Supplementary Table S1-4

### Supplementary Figures

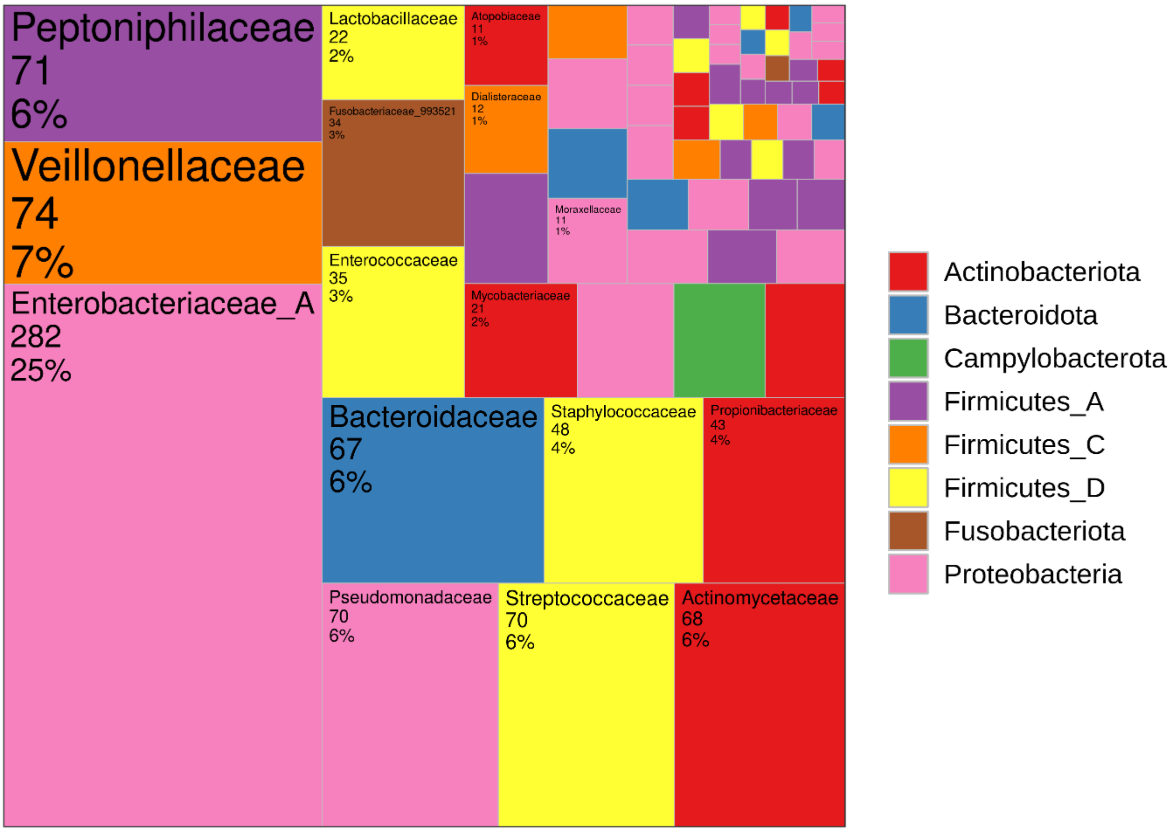


**Supplementary Fig. S1. Treemap showing the fraction of ASVs assigned to Family.**

The number and percentages of ASVs belong to each family are denoted inside of each box. The colors depict corresponding phyla.

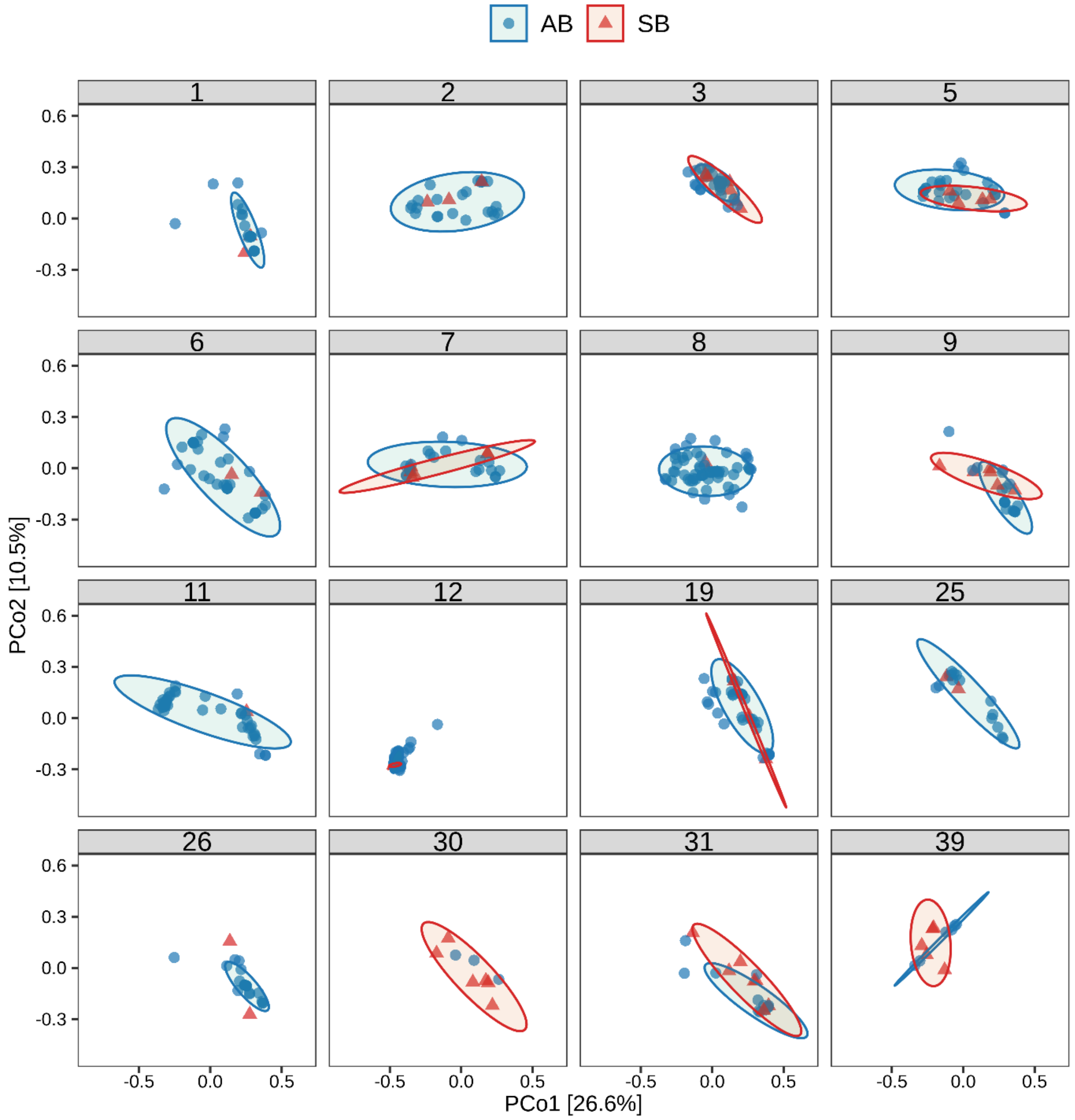


**Supplementary Fig. S2. Beta diversity (unweighted unifrac) analysis in participants with at least one symptomatic bacteriuria event.**

Principal coordinates analyses (PCoA) of beta-diversity between asymptomatic (AB, blue) and symptomatic (SB, red) bacteriuria group based on unweighted unifrac distance matrices. Each dot represents an individual sample.

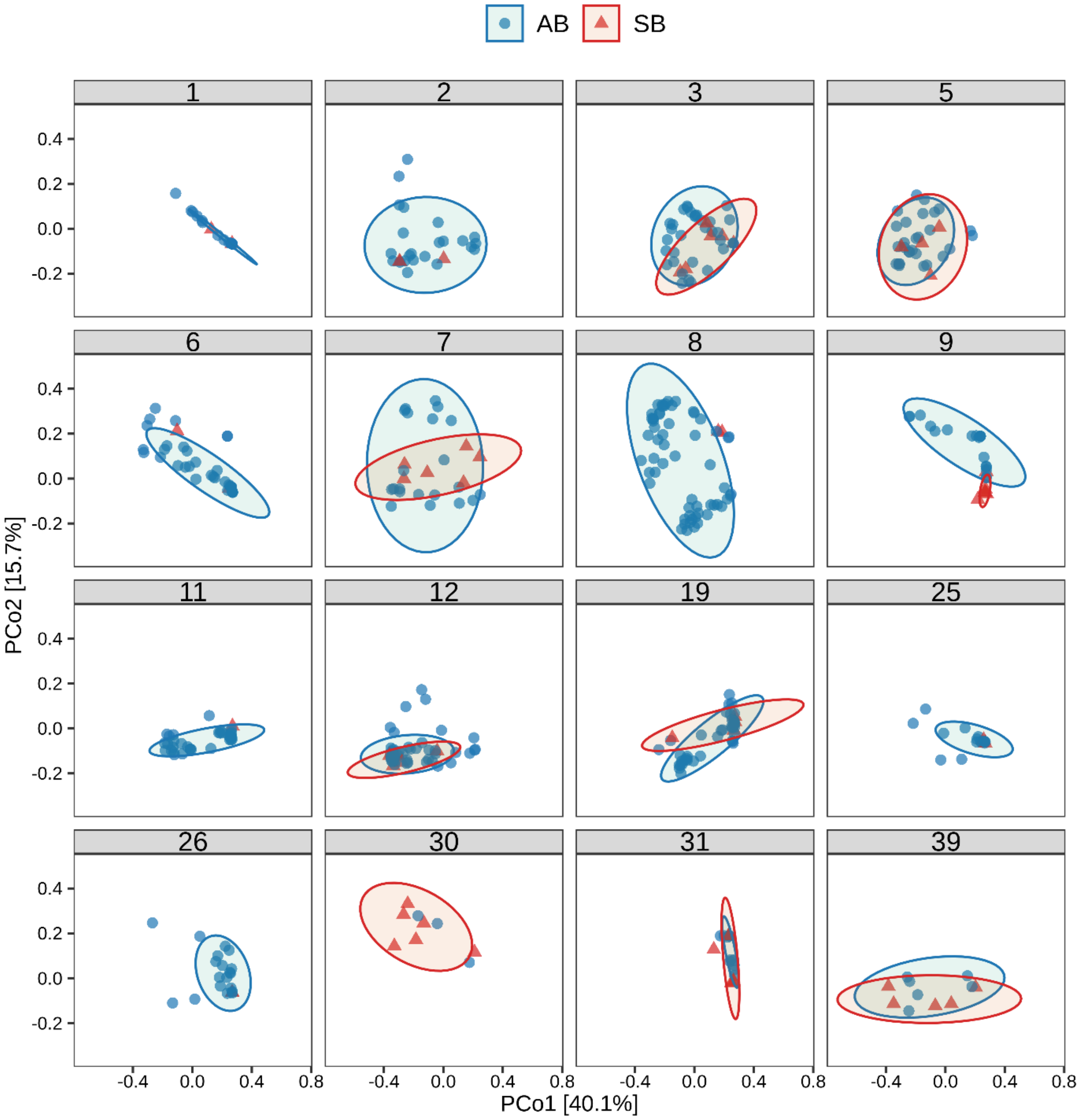


**Supplementary Fig. S3. Beta diversity analysis (weighted unifrac) in participants with at least one symptomatic bacteriuria event.**

Principal coordinates analyses (PCoA) of beta-diversity between asymptomatic (AB, blue) and symptomatic (SB, red) bacteriuria group based on weighted unifrac distance matrices. Each dot represents an individual sample.

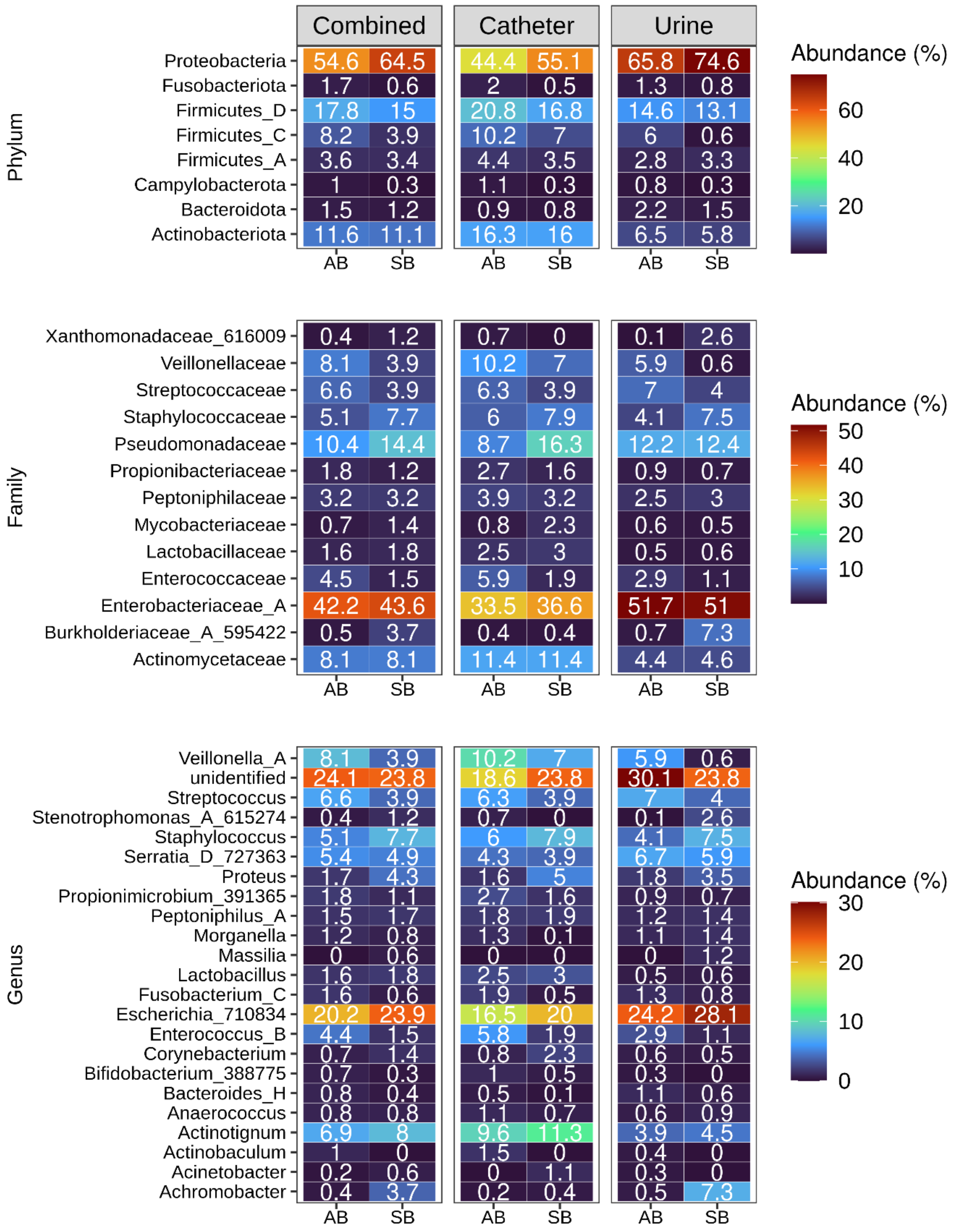


**Supplementary Fig. S4. Overview of taxonomic composition in asymptomatic and symptomatic bacteriuria groups across combined, catheter and urine dataset.** The colour heatmaps depicting mean relative abundances in percentages ranging from blue (low abundance) to red (high abundance) grouped by phylum, family, and genus. The numbers inside heatmaps show mean relative abundance of corresponding taxa indicated in y-axis across asymptomatic bacteriuria (AB) and symptomatic (SB) groups. Genus are shown if their mean relative abundances in any of the group was more than 1.

### Supplementary Tables

**Supplementary Table S1.** Participants information and samples analysed.

| Participant | Received | Analysed | 18-months completed |
|---|---|---|---|
| 1 | 10 | 10 | Deceased |
| 2 | 15 | 15 | Yes |
| 3 | 20 | 20 | Yes |
| 4 | 5 | 5 | Deceased |
| 5 | 17 | 17 | Yes |
| 6 | 20 | 20 | Yes |
| 7 | 22 | 15 | Yes |
| 8 | 32 | 32 | Yes |
| 9 | 15 | 13 | Yes |
| 10 | Interviewed but excluded because of the criteria did not met. | | |
| 11 | 24 | 20 | Yes |
| 12 | 31 | 31 | Yes |
| 13 | 19 | 16 | Yes |
| 14 | 6 | 6 | Deceased |
| 15 | Interviewed but excluded because of the criteria did not met. | | |
| 16 | 21 | 17 | Yes |
| 17 | Interviewed but excluded because of the criteria did not met. | | |
| 18 | Interviewed but excluded because of the criteria did not met. | | |
| 19 | 26 | 21 | Yes |
| 20 | Interviewed but excluded because of the criteria did not met. | | |
| 21 | Interviewed but excluded because of the criteria did not met. | | |
| 22 | Interviewed but excluded because of the criteria did not met. | | |
| 23 | Interviewed but excluded because of the criteria did not met. | | |
| 24 | 8 | 5 | No |
| 25 | 16 | 9 | No |
| 26 | 13 | 11 | No |
| 27 | 1 | 1 | withdrawn |
| 28 | Interviewed but excluded because of the criteria did not met. | | |
| 29 | 7 | 5 | No |
| 30 | 6 | 5 | No |
| 31 | 14 | 10 | No |
| 32 | Interviewed but excluded because of the criteria did not met. | | |
| 33 | 7 | 1 | No |
| 34 | 2 | 1 | No |
| 35 | 5 | 1 | No |
| 36 | Interviewed but excluded because of the criteria did not met. | | |
| 37 | Interviewed but excluded because of the criteria did not met. | | |
| 38 | 4 | 2 | No |
| 39 | 13 | 6 | No |

**Supplementary Table S2.** Machine learning models.

| ID | Name | Reference |
|---|---|---|
| lr | Logistic Regression | sklearn.linear_model._logistic.LogisticRegression |
| knn | K Neighbors Classifier | sklearn.neighbors._classification.KNeighborsClassifier |
| nb | Naive Bayes | sklearn.naive_bayes.GaussianNB |
| dt | Decision Tree Classifier | sklearn.tree._classes.DecisionTreeClassifier |
| svm | SVM - Linear Kernel | sklearn.linear_model._stochastic_gradient.SGDClassifier |
| rbfsvm | SVM - Radial Kernel | sklearn.svm._classes.SVC |
| gpc | Gaussian Process Classifier | sklearn.gaussian_process._gpc.GaussianProcessClassifier |
| mlp | MLP Classifier | sklearn.neural_network._multilayer_perceptron.MLPClassifier |
| ridge | Ridge Classifier | sklearn.linear_model._ridge.RidgeClassifier |
| rf | Random Forest Classifier | sklearn.ensemble._forest.RandomForestClassifier |
| qda | Quadratic Discriminant Analysis | sklearn.discriminant_analysis.QuadraticDiscriminantAnalysis |
| ada | Ada Boost Classifier | sklearn.ensemble._weight_boosting.AdaBoostClassifier |
| gbc | Gradient Boosting Classifier | sklearn.ensemble._gb.GradientBoostingClassifier |
| lda | Linear Discriminant Analysis | sklearn.discriminant_analysis.LinearDiscriminantAnalysis |
| et | Extra Trees Classifier | sklearn.ensemble._forest.ExtraTreesClassifier |
| xgboost | Extreme Gradient Boosting | xgboost.sklearn.XGBClassifier |
| lightgbm | Light Gradient Boosting Machine | lightgbm.sklearn.LGBMClassifier |
| catboost | CatBoost Classifier | catboost.core.CatBoostClassifier |
| dummy | Dummy Classifier | sklearn.dummy.DummyClassifier |

**Supplementary Table S3** Performance of machine learning models during cross-validation.

| | | Cross-validation set | | | | | | | | | | | |
|---|---|---|---|---|---|---|---|---|---|---|---|---|---|
| | | Without antibiotic treated samples | | | | | | With antibiotic treated samples | | | | | |
| | | Combined | | Catheter | | Urine | | Combined | | Catheter | | Urine | |
| | | ASV | Taxa | ASV | Taxa | ASV | Taxa | ASV | Taxa | ASV | Taxa | ASV | Taxa |
| AUROC | min | 0.91 | 0.78 | 0.83 | 0.72 | 0.87 | 0.72 | 0.82 | 0.72 | 0.73 | 0.67 | 0.68 | 0.67 |
| AUROC | max | 0.98 | 0.91 | 0.98 | 0.97 | 0.98 | 0.87 | 0.89 | 0.84 | 0.91 | 0.82 | 0.84 | 0.8 |
| AUROC | q25 | 0.93 | 0.86 | 0.88 | 0.82 | 0.92 | 0.76 | 0.83 | 0.77 | 0.81 | 0.72 | 0.73 | 0.71 |
| AUROC | q75 | 0.96 | 0.88 | 0.93 | 0.87 | 0.95 | 0.83 | 0.86 | 0.81 | 0.86 | 0.77 | 0.78 | 0.75 |
| AUROC | mean | 0.95 | 0.87 | 0.9 | 0.84 | 0.94 | 0.8 | 0.85 | 0.79 | 0.83 | 0.74 | 0.76 | 0.73 |
| AUROC | std | 0.02 | 0.03 | 0.04 | 0.06 | 0.03 | 0.04 | 0.02 | 0.03 | 0.05 | 0.04 | 0.04 | 0.04 |
| Precision | min | 0.94 | 0.94 | 0.93 | 0.91 | 0.94 | 0.91 | 0.9 | 0.85 | 0.1 | 0.85 | 0.82 | 0.84 |
| Precision | max | 0.97 | 0.96 | 0.97 | 0.96 | 0.97 | 0.95 | 0.91 | 0.9 | 0.91 | 0.89 | 0.9 | 0.89 |
| Precision | q25 | 0.95 | 0.94 | 0.94 | 0.93 | 0.95 | 0.93 | 0.91 | 0.88 | 0.89 | 0.86 | 0.88 | 0.87 |
| Precision | q75 | 0.96 | 0.95 | 0.95 | 0.95 | 0.96 | 0.94 | 0.91 | 0.9 | 0.9 | 0.88 | 0.89 | 0.88 |
| Precision | mean | 0.95 | 0.95 | 0.95 | 0.94 | 0.95 | 0.93 | 0.91 | 0.89 | 0.84 | 0.87 | 0.88 | 0.87 |
| Precision | std | 0.01 | 0.01 | 0.01 | 0.01 | 0.01 | 0.01 | 0 | 0.01 | 0.19 | 0.01 | 0.02 | 0.01 |
| Recall | min | 0.76 | 0.33 | 0.59 | 0.61 | 0.47 | 0.45 | 0.39 | 0.4 | 0.11 | 0.5 | 0.17 | 0.48 |
| Recall | max | 0.97 | 0.96 | 0.94 | 0.93 | 0.9 | 0.95 | 0.64 | 0.86 | 0.86 | 0.84 | 0.7 | 0.84 |
| Recall | q25 | 0.83 | 0.58 | 0.77 | 0.7 | 0.74 | 0.62 | 0.42 | 0.53 | 0.68 | 0.61 | 0.43 | 0.55 |
| Recall | q75 | 0.91 | 0.78 | 0.83 | 0.88 | 0.84 | 0.86 | 0.6 | 0.66 | 0.77 | 0.71 | 0.59 | 0.62 |
| Recall | mean | 0.87 | 0.68 | 0.8 | 0.79 | 0.78 | 0.75 | 0.52 | 0.6 | 0.68 | 0.66 | 0.51 | 0.6 |
| Recall | std | 0.06 | 0.18 | 0.07 | 0.11 | 0.1 | 0.15 | 0.1 | 0.12 | 0.2 | 0.09 | 0.12 | 0.09 |
| F1 | min | 0.82 | 0.43 | 0.69 | 0.71 | 0.58 | 0.56 | 0.45 | 0.47 | 0.04 | 0.58 | 0.14 | 0.56 |
| F1 | max | 0.96 | 0.95 | 0.94 | 0.93 | 0.92 | 0.94 | 0.71 | 0.85 | 0.87 | 0.85 | 0.76 | 0.84 |
| F1 | q25 | 0.87 | 0.68 | 0.83 | 0.78 | 0.81 | 0.71 | 0.5 | 0.6 | 0.74 | 0.68 | 0.5 | 0.62 |
| F1 | q75 | 0.93 | 0.83 | 0.87 | 0.9 | 0.88 | 0.89 | 0.67 | 0.72 | 0.81 | 0.76 | 0.67 | 0.69 |
| F1 | mean | 0.9 | 0.75 | 0.85 | 0.84 | 0.83 | 0.81 | 0.59 | 0.67 | 0.71 | 0.71 | 0.57 | 0.67 |
| F1 | std | 0.04 | 0.15 | 0.05 | 0.07 | 0.08 | 0.11 | 0.1 | 0.1 | 0.22 | 0.07 | 0.14 | 0.07 |
| Accuracy | min | 0.76 | 0.33 | 0.59 | 0.61 | 0.47 | 0.45 | 0.39 | 0.4 | 0.11 | 0.5 | 0.17 | 0.48 |
| Accuracy | max | 0.97 | 0.96 | 0.94 | 0.93 | 0.9 | 0.95 | 0.64 | 0.86 | 0.86 | 0.84 | 0.7 | 0.84 |
| Accuracy | q25 | 0.83 | 0.58 | 0.77 | 0.7 | 0.74 | 0.62 | 0.42 | 0.53 | 0.68 | 0.61 | 0.43 | 0.55 |
| Accuracy | q75 | 0.91 | 0.78 | 0.83 | 0.88 | 0.84 | 0.86 | 0.6 | 0.66 | 0.77 | 0.71 | 0.59 | 0.62 |
| Accuracy | mean | 0.87 | 0.68 | 0.8 | 0.79 | 0.78 | 0.75 | 0.52 | 0.6 | 0.68 | 0.66 | 0.51 | 0.6 |
| Accuracy | std | 0.06 | 0.18 | 0.07 | 0.11 | 0.1 | 0.15 | 0.1 | 0.12 | 0.2 | 0.09 | 0.12 | 0.09 |
| Balanced accuracy | min | 0.72 | 0.65 | 0.63 | 0.58 | 0.72 | 0.59 | 0.66 | 0.61 | 0.5 | 0.61 | 0.54 | 0.58 |
| Balanced accuracy | max | 0.89 | 0.82 | 0.89 | 0.88 | 0.93 | 0.82 | 0.77 | 0.78 | 0.82 | 0.72 | 0.75 | 0.72 |
| Balanced accuracy | q25 | 0.83 | 0.7 | 0.78 | 0.66 | 0.82 | 0.66 | 0.67 | 0.67 | 0.74 | 0.64 | 0.63 | 0.63 |
| Balanced accuracy | q75 | 0.86 | 0.79 | 0.83 | 0.78 | 0.88 | 0.75 | 0.75 | 0.72 | 0.78 | 0.7 | 0.7 | 0.69 |
| Balanced accuracy | mean | 0.84 | 0.75 | 0.8 | 0.72 | 0.84 | 0.71 | 0.72 | 0.69 | 0.74 | 0.67 | 0.66 | 0.66 |
| Balanced accuracy | std | 0.04 | 0.05 | 0.07 | 0.09 | 0.05 | 0.07 | 0.04 | 0.04 | 0.08 | 0.04 | 0.05 | 0.04 |

**Supplementary Table S4.** Performance of machine learning models on held-out set.

| | | Held-out set | | | | | | | | | | | |
|---|---|---|---|---|---|---|---|---|---|---|---|---|---|
| | | Without antibiotic treated samples | | | | | | With antibiotic treated samples | | | | | |
| | | Combined | | Catheter | | Urine | | Combined | | Catheter | | Urine | |
| | | ASV | Taxa | ASV | Taxa | ASV | Taxa | ASV | Taxa | ASV | Taxa | ASV | Taxa |
| AUROC | min | 0.85 | 0.69 | 0.7 | 0.68 | 0.85 | 0.57 | 0.69 | 0.55 | 0.54 | 0.53 | 0.47 | 0.48 |
| AUROC | max | 1 | 0.99 | 0.99 | 0.96 | 1 | 1 | 0.93 | 0.86 | 0.96 | 0.85 | 0.84 | 0.8 |
| AUROC | q25 | 0.93 | 0.78 | 0.85 | 0.76 | 0.93 | 0.69 | 0.81 | 0.66 | 0.74 | 0.61 | 0.6 | 0.58 |
| AUROC | q75 | 0.98 | 0.88 | 0.96 | 0.87 | 0.98 | 0.89 | 0.88 | 0.76 | 0.9 | 0.69 | 0.74 | 0.75 |
| AUROC | mean | 0.95 | 0.83 | 0.88 | 0.81 | 0.95 | 0.79 | 0.83 | 0.71 | 0.82 | 0.66 | 0.67 | 0.65 |
| AUROC | std | 0.04 | 0.08 | 0.1 | 0.08 | 0.03 | 0.13 | 0.07 | 0.08 | 0.11 | 0.09 | 0.11 | 0.11 |
| Precision | min | 0.93 | 0.91 | 0.92 | 0.87 | 0.94 | 0.89 | 0.87 | 0.81 | 0.86 | 0.8 | 0.81 | 0.78 |
| Precision | max | 0.97 | 0.96 | 0.96 | 0.98 | 0.98 | 0.97 | 0.92 | 0.92 | 0.93 | 0.92 | 0.92 | 0.92 |
| Precision | q25 | 0.95 | 0.93 | 0.94 | 0.91 | 0.96 | 0.92 | 0.91 | 0.86 | 0.9 | 0.84 | 0.86 | 0.85 |
| Precision | q75 | 0.96 | 0.95 | 0.95 | 0.95 | 0.97 | 0.96 | 0.91 | 0.89 | 0.93 | 0.88 | 0.91 | 0.88 |
| Precision | mean | 0.96 | 0.94 | 0.95 | 0.92 | 0.96 | 0.94 | 0.91 | 0.88 | 0.91 | 0.86 | 0.87 | 0.86 |
| Precision | std | 0.01 | 0.01 | 0.01 | 0.03 | 0.01 | 0.02 | 0.01 | 0.03 | 0.02 | 0.03 | 0.03 | 0.04 |
| Recall | min | 0.73 | 0.32 | 0.63 | 0.65 | 0.54 | 0.5 | 0.35 | 0.36 | 0.11 | 0.44 | 0.25 | 0.39 |
| Recall | max | 0.97 | 0.93 | 0.96 | 0.98 | 0.96 | 0.93 | 0.68 | 0.88 | 0.9 | 0.82 | 0.68 | 0.82 |
| Recall | q25 | 0.82 | 0.55 | 0.77 | 0.71 | 0.73 | 0.6 | 0.43 | 0.53 | 0.65 | 0.53 | 0.39 | 0.53 |
| Recall | q75 | 0.92 | 0.75 | 0.92 | 0.88 | 0.88 | 0.87 | 0.62 | 0.62 | 0.78 | 0.73 | 0.58 | 0.63 |
| Recall | mean | 0.87 | 0.67 | 0.82 | 0.8 | 0.8 | 0.76 | 0.53 | 0.58 | 0.67 | 0.63 | 0.48 | 0.58 |
| Recall | std | 0.06 | 0.16 | 0.09 | 0.1 | 0.1 | 0.14 | 0.11 | 0.11 | 0.21 | 0.12 | 0.11 | 0.1 |
| F1 | min | 0.8 | 0.42 | 0.73 | 0.74 | 0.66 | 0.62 | 0.42 | 0.43 | 0.05 | 0.53 | 0.27 | 0.46 |
| F1 | max | 0.97 | 0.92 | 0.95 | 0.98 | 0.96 | 0.94 | 0.74 | 0.87 | 0.91 | 0.85 | 0.75 | 0.84 |
| F1 | q25 | 0.87 | 0.66 | 0.83 | 0.79 | 0.81 | 0.72 | 0.51 | 0.61 | 0.72 | 0.63 | 0.46 | 0.61 |
| F1 | q75 | 0.93 | 0.82 | 0.92 | 0.9 | 0.91 | 0.9 | 0.7 | 0.69 | 0.82 | 0.78 | 0.66 | 0.7 |
| F1 | mean | 0.9 | 0.75 | 0.86 | 0.84 | 0.85 | 0.82 | 0.61 | 0.66 | 0.71 | 0.7 | 0.55 | 0.65 |
| F1 | std | 0.04 | 0.13 | 0.06 | 0.07 | 0.07 | 0.1 | 0.11 | 0.09 | 0.23 | 0.09 | 0.12 | 0.09 |
| Accuracy | min | 0.73 | 0.32 | 0.63 | 0.65 | 0.54 | 0.5 | 0.35 | 0.36 | 0.11 | 0.44 | 0.25 | 0.39 |
| Accuracy | max | 0.97 | 0.93 | 0.96 | 0.98 | 0.96 | 0.93 | 0.68 | 0.88 | 0.9 | 0.82 | 0.68 | 0.82 |
| Accuracy | q25 | 0.82 | 0.55 | 0.77 | 0.71 | 0.73 | 0.6 | 0.43 | 0.53 | 0.65 | 0.53 | 0.39 | 0.53 |
| Accuracy | q75 | 0.92 | 0.75 | 0.92 | 0.88 | 0.88 | 0.87 | 0.62 | 0.62 | 0.78 | 0.73 | 0.58 | 0.63 |
| Accuracy | mean | 0.87 | 0.67 | 0.82 | 0.8 | 0.8 | 0.76 | 0.53 | 0.58 | 0.67 | 0.63 | 0.48 | 0.58 |
| Accuracy | std | 0.06 | 0.16 | 0.09 | 0.1 | 0.1 | 0.14 | 0.11 | 0.11 | 0.21 | 0.12 | 0.11 | 0.1 |
| Balanced_accuracy | min | 0.7 | 0.57 | 0.64 | 0.45 | 0.67 | 0.32 | 0.64 | 0.46 | 0.51 | 0.42 | 0.49 | 0.44 |
| Balanced_accuracy | max | 0.97 | 0.86 | 0.96 | 0.88 | 0.98 | 0.91 | 0.82 | 0.75 | 0.88 | 0.75 | 0.78 | 0.81 |
| Balanced_accuracy | q25 | 0.81 | 0.65 | 0.73 | 0.63 | 0.85 | 0.53 | 0.68 | 0.62 | 0.67 | 0.56 | 0.58 | 0.58 |
| Balanced_accuracy | q75 | 0.92 | 0.75 | 0.89 | 0.82 | 0.93 | 0.78 | 0.77 | 0.7 | 0.83 | 0.67 | 0.67 | 0.68 |
| Balanced_accuracy | mean | 0.86 | 0.71 | 0.82 | 0.71 | 0.88 | 0.66 | 0.72 | 0.65 | 0.74 | 0.61 | 0.63 | 0.63 |
| Balanced_accuracy | std | 0.08 | 0.08 | 0.09 | 0.13 | 0.07 | 0.16 | 0.05 | 0.07 | 0.11 | 0.09 | 0.07 | 0.1 |
| AUPRC | min | 0.37 | 0.13 | 0.13 | 0.16 | 0.18 | 0.07 | 0.21 | 0.14 | 0.14 | 0.12 | 0.11 | 0.11 |
| AUPRC | max | 1 | 0.85 | 0.87 | 0.74 | 1 | 1 | 0.62 | 0.51 | 0.82 | 0.32 | 0.53 | 0.54 |
| AUPRC | q25 | 0.6 | 0.26 | 0.24 | 0.2 | 0.35 | 0.14 | 0.34 | 0.19 | 0.29 | 0.14 | 0.16 | 0.14 |
| AUPRC | q75 | 0.84 | 0.48 | 0.6 | 0.47 | 0.58 | 0.39 | 0.53 | 0.33 | 0.62 | 0.22 | 0.3 | 0.33 |
| AUPRC | mean | 0.7 | 0.39 | 0.45 | 0.36 | 0.49 | 0.3 | 0.43 | 0.27 | 0.48 | 0.19 | 0.25 | 0.25 |
| AUPRC | std | 0.18 | 0.2 | 0.22 | 0.19 | 0.2 | 0.25 | 0.12 | 0.1 | 0.21 | 0.07 | 0.11 | 0.13 |
| Baseline | | 0.05 | 0.05 | 0.06 | 0.06 | 0.04 | 0.04 | 0.1 | 0.1 | 0.1 | 0.1 | 0.11 | 0.11 |
